# Old age variably impacts chimpanzee engagement and efficiency in stone tool use

**DOI:** 10.1101/2024.11.25.625128

**Authors:** E Howard-Spink, T Matsuzawa, S Carvalho, C Hobaiter, K Almeida-Warren, T Gruber, D Biro

## Abstract

We know vanishingly little about how long-lived apes experience senescence in the wild, particularly with respect to their foraging behaviors, which are essential for survival. Some wild apes use tools during foraging and, given the additional cognitive and physical challenges presented by tool use, we predict that such behaviors are at a heightened risk of senescence. However, until the present, longitudinal analysis of the effects of progressive aging on wild ape tool-use behaviors has not been possible due to a lack of available data. In response to this research gap, we sampled data from a longitudinal video archive that contained footage of wild chimpanzees engaging in one of their most complex forms of tool use - the cracking of hard-shelled nuts with hammers and anvil stones, termed *nut cracking* - at an ‘outdoor laboratory’ at Bossou, Guinea. By sampling data over a 17-year period, we describe how the extent to which wild chimpanzees engage in – and efficiently perform – nut cracking changes between the ages of approximately 39-44 to 56-61 years of age. Over this extended sampling period, chimpanzees began attending experimental nut cracking sites less frequently than younger individuals. Several elderly chimpanzees exhibited reductions in efficiency across multiple components of nut cracking, including taking more time to select stone tools prior to use, and taking longer to crack open nuts and consume the associated pieces of kernel. Two chimpanzees also began using less streamlined behavioral sequences to crack nuts, including a greater number of actions (such as more numerous strikes of the hammer stone). Most notably, we report interindividual variability in the extent to which elderly chimpanzees’ tool-use behaviors changed during our sample period – ranging from small to profound reductions in tool engagement and efficiency – as well as differences in the specific aspects of nut cracking behaviors that changed for each individual as they aged. We discuss the possible causes of these changes with reference to research into senescence in captive primates, and provide future directions for research of primate aging in both captive and wild settings.

## 1. Introduction

Senescence can be understood as a reduction in the efficacy of biological systems with increasing age. In recent decades, the senescence of non-human primates (henceforth primates) have been a point of considerable research interest^1–3^. In part, this is due to the fact that primates offer uniquely translatable models for understanding the nature and evolution of human senescence, given their high genetic, physiological, and behavioral similarities^1,4–6^. Accordingly, many parallels between the senescent processes of humans and primates have been uncovered, including in their cognition^7–12^ physiology^1,13–15^, and their resultant behaviours^16–20^. However, thus far, the majority of these studies have focused on captive populations of short-living species, and exploit existing variation in age among individuals to infer the effects of senescence through cross-sectional analyses^1,8,21,22^. Comparatively few studies have investigated how senescence influences behaviors of primates in the wild (with the exception of a handful of examples^23–27)^, and even fewer employ longitudinal data sampling to understand how senescence leads to changes in the behaviors of specific individuals over time. These studies of wild individuals are highly valuable as they shed light on how aging affects the day-to-day behaviors that primates rely on for survival – conclusions which cannot be easily predicted from studies from captivity, given that patterns of physiological senescence can differ significantly between captive and naturalistic environments^28,29^. Further research is therefore required to understand how aging influences the lives of wild primates. This is particularly true for long-living great ape species, which are underrepresented in the existing literature, yet their phylogenetic proximity to humans means that they likely offer the most suitable models for understanding the evolution of the human aging process.

Of the day-to-day behaviors of great apes, the combined physical and cognitive demands of tool use make it likely to exhibit particular changes with progressive old age. The tool-use behaviors of great apes are highly challenging, requiring tool users to draw upon a wide array of high-level cognitive abilities, including planning and the flexible assembly of actions into goal-directed behaviors^30–33;^ an understanding of causal relationships between objects^34–36;^ as well as knowledge of the physical properties of objects and how to exploit these properties to use tools successfully^32,37–39^. Many of these cognitive abilities – including the more specific subcomponents of these abilities, such as motor coordination^12^, working memory^40^, executive functioning^10^, and cognitive flexibility^11^ – have been identified as at risk of senescence in captive living primates; however, never in tool-use behaviors themselves. Moreover, physiological changes including reduced bone mass^13,15,41^, muscle wasting (sarcopenia^15,42,43^), the development of arthritis^15^, and reduced visual acuity^23^ have been identified in a number of primate species, including great apes, which all influence individual strength, dexterity, and accuracy of movement. These findings suggest that tool use could be a domain of great-ape behavior that is at specific risk of senescence; however, it has not yet been possible to address this question due to an absence of long-term data on great ape tool-use behaviors that also include instances of tool use performed by elderly individuals.

To address this gap in the literature, we used a longitudinal archive of video footage of wild, West African chimpanzees (*Pan troglodytes verus*) using stone tools to crack hard-shelled nuts as part of a decades-long field experiment in a clearing in the Bossou forest, Guinea^44,45^ (’the outdoor laboratory’; see Methods section 4.1 and Fig. 1 for information about the experimental set up at the outdoor laboratory). Nut cracking is one of the most sophisticated forms of tool use observed in the animal kingdom, and is habitually performed by a number of primate species in the wild including several communities of chimpanzees^37,46,47^ (*Pan troglodytes*), capuchin monkeys^48–52^ (*Cebus* and *Sapajus* spp.), and long-tailed macaques^53,54^ (*Macaca fascicularis*). During nut cracking, a nut is placed on a hard substrate (for the chimpanzee population used in this study, this substrate is a portable anvil stone, though at other sites this may be a root or rocky outcrop^37,39^), before being struck repeatedly with a hammer stone until the nut is cracked open and the edible kernel inside extracted and consumed (see Fig. 1a). Nut cracking is learnt by chimpanzees at Bossou by the age of seven years (often between 3.5 and 5 years of age^55,56^), and chimpanzees continue to perform this behavior throughout their lives^44^. This life-long engagement in nut cracking permits characterization of how individual chimpanzees perform this behavior across their lives^57^, including during the oldest years of their lifespan. Nut cracking features all of the key elements of tool-use behaviors which we predict will make them likely to senesce with old age, including the need to construct complex object associations^34,35,58^; to organize and address multiple goals using an extended behavioral sequence^32,33^; to select tool objects based on perceived properties^32,37,39^, and to combine objects with sufficient dexterity, precision and force and to crack nuts without knocking nuts off the anvil, or smashing interior kernels^38^.

**Figure 1.**
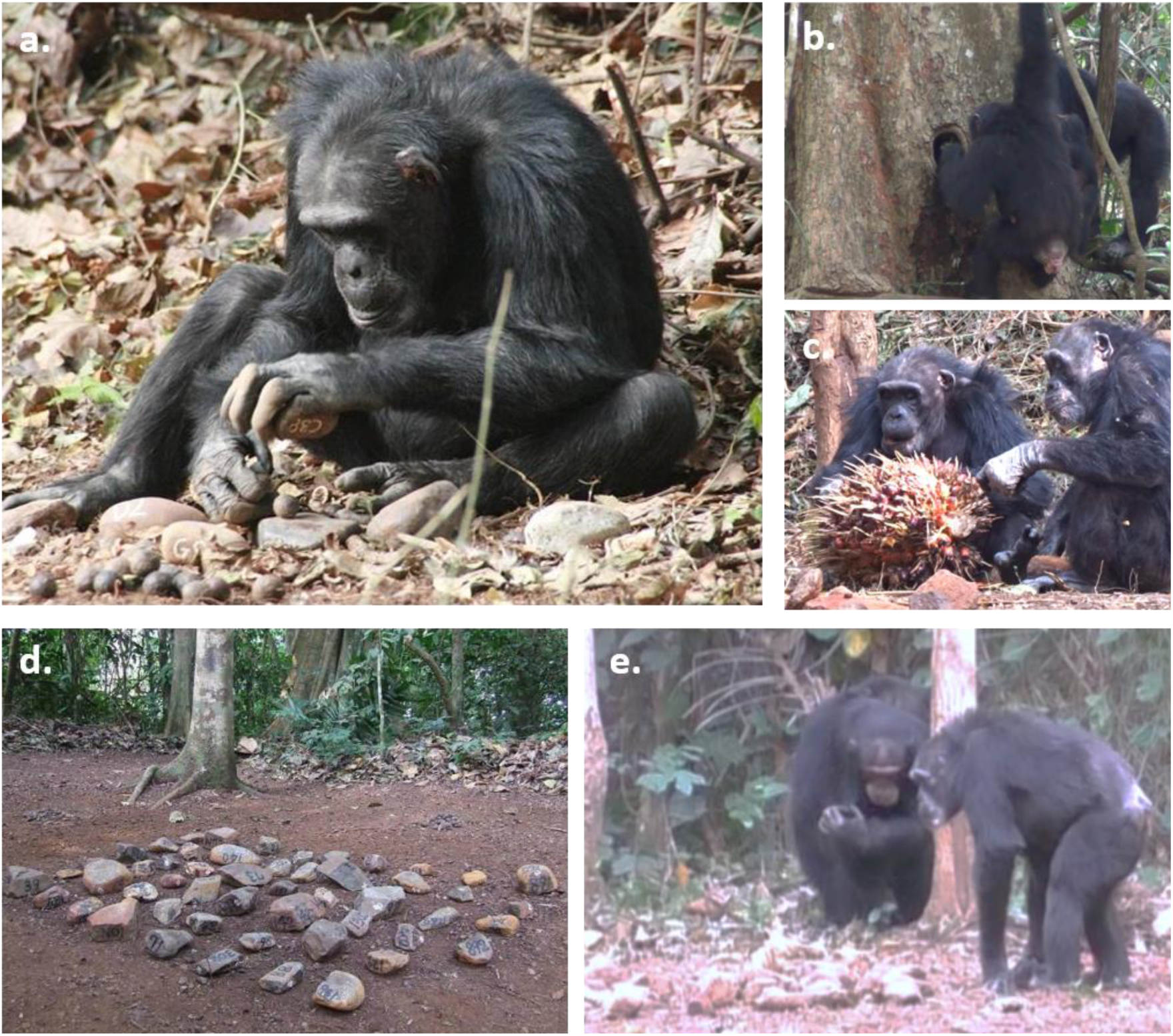
The experimental set-up and behaviors at the outdoor laboratory. (a) Yo, an elderly female (approx. 51 years old), has placed a coula nut (*Coula edulis*) on an anvil stone, and is preparing to strike with the hammer stone. (b) An adult female (14 years old) using a leaf tool to drink water from the water point. (c) Yo and Velu (left to right, both approximately 56-58 years old), eating oil-palm fruits from an available raceme. (d) The central stone-tool matrix with numbered stones. (e) Yo (right; approximately 49 years old) selecting tools from the stone matrix.

To determine whether – and if so, how – old age influenced the nut cracking behaviors of wild chimpanzees, we sampled video footage collected at the outdoor laboratory in five field seasons spanning a 17-year timeframe between 1999 and 2016. During this timeframe, we collected longitudinal data on five chimpanzees as they aged from approximately 39-44 years old (end of mid-life) to 56-61 years old (ages which are close to the maximum for wild chimpanzee lifespans^59^). Across sampled field seasons, we estimated how readily chimpanzees engaged in nut cracking behaviors using two key metrics: the frequency which chimpanzees visited outdoor laboratory sites during each field season (compared to a control group of younger individuals between the ages of 8 and 30 years of age), and when present, the proportion of time chimpanzees spent engaging with nuts and stone tools compared to other key behaviors (such as drinking water from an available water point using leaf tools^60^ and eating oil-palm fruits that are provisioned alongside nuts; see Figs 1b & 1c). We also measured how quickly chimpanzees selected stone tools from a central matrix of over 50 stones (a generalized metric of efficiency during tool selection; see Figs 1d & 1e), as well as how efficiently chimpanzees cracked open nuts using stone tools and consumed the associated kernel.

Overall, with increasingly old age (measured by progressive field seasons), elderly chimpanzees were significantly less likely to visit experimental nut cracking sites. In comparison, younger individuals (i.e. younger adults, and older immatures who had learnt how to crack nuts) exhibited no change over successive field seasons, suggesting that this reduced attendance at experimental nut cracking sites was confined to an elderly, aging cohort of chimpanzees. When present at the outdoor laboratory, two of our five elderly individuals spent substantially less time engaging in nut cracking at older ages, as compared to their behaviors in previous field seasons. Moreover, several elderly chimpanzees took more time to select stone tools in later field seasons than in earlier field seasons, and were also less efficient at using stone tools to crack open oil-palm nuts. Importantly, we detected considerable interindividual variability in the effects of aging on how readily elderly chimpanzees engaged with nuts and stone tools, how quickly they selected tools, and the efficiency with which elderly chimpanzees used tools. Some individuals demonstrated little-to-no change in engagement and efficiency with aging, whereas others experienced significant changes across different aspects of their nut cracking behavior. We discuss our findings in the context of existing literature for primate aging, and hypothesize which factors underpin observed changes in behavior, including cognitive, physiological, and motivational causes, and generate a number of suggested avenues for further study. We also discuss how aging can likely compound existing interindividual differences in the proficiency of socially-learnt technical skills in non-human animals, and discuss how tool use could present a suitable indicator for identifying individuals who are experiencing the effects of profound senescence in the wild.

## 2. Results

### 2.1 Sampled data

We sampled five timepoints separated by intervals of 3-5 years (field seasons 1999-2000, 2004-2005, 2008-2009, 2011-2012, and 2016-2017; with each field season referred to hereafter by its initial year). The exact age at which chimpanzees may be considered ‘old’ is a point of continued debate. However, chimpanzees are generally considered to begin entering old age at approximately 40 years old^43,61,62^, after which survivorship begins to decrease more rapidly than earlier in adulthood^59^. We therefore confined our analysis to individuals who were above at least 30 years of age in at least three of the five sampled field seasons. This sample allowed us to collect data longitudinally, beginning with ages where chimpanzees are in the prime of adulthood, and spanning into old age. Under these criteria, we were able to include four old-age females (Fana, Jire, Velu and Yo) and one old-age male (Tua) in our analyses. All five of these individuals were present at Bossou when the long-term research project was established in 1976; as a consequence, their birth years are estimates based on physical growth characteristics at the time of first observation (see Table 1). Given that these estimated birth years range from 1956 to 1961, these individuals are estimated to be between 39-44 years old at the start of our sampling window (1999), and therefore are entering the window of ‘old age’. By 2016, estimated ages ranged from 56-61 years old, reflecting an age which is near the maximum for wild chimpanzees. All four females were present at Bossou throughout the entire sampled time-frame for our analysis; however, Tua, the only adult male, disappeared in September 2013 (presumed dead). Therefore, data for Tua spans a shorter time frame between the 1999 and 2011 field seasons. As all five focal individuals’ ages are within five years of each other (and therefore similar estimates), we use the progression through field seasons as a proxy for increasing age, and treat all individuals as similarly ‘old’ at each field season.

**Table 1.**
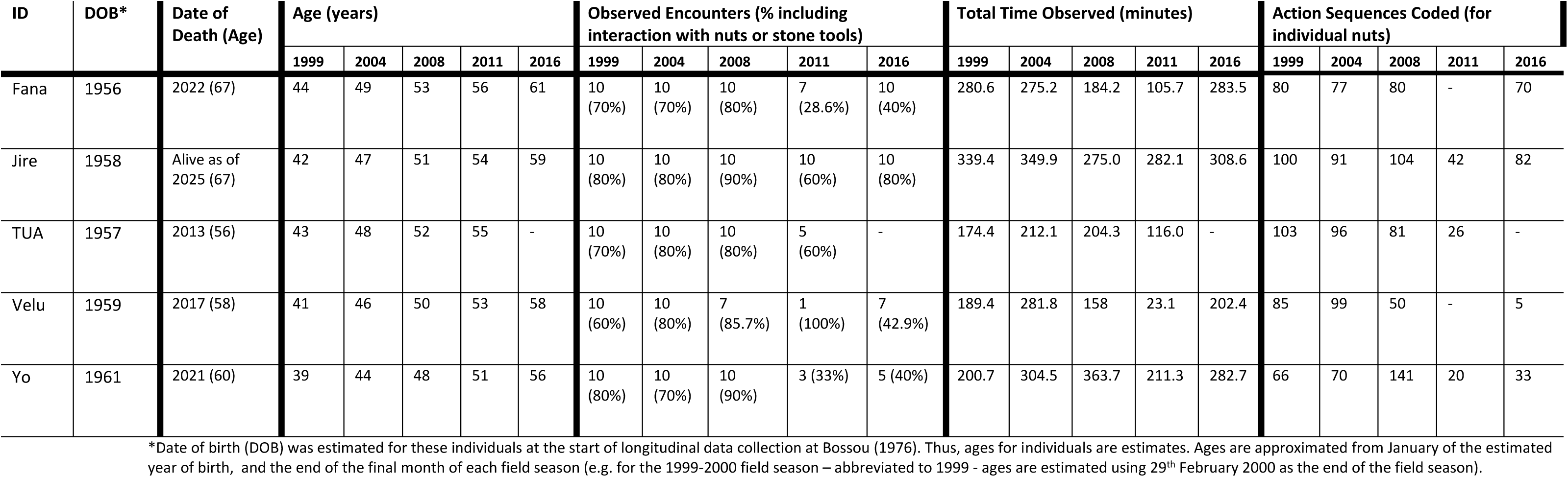
Summary of sampled observations for each focal old-age individual. Total time observed includes all time individuals were present in the first 10 encounters of each field season (Observed Encounters). Dashed lines (-) represent where no data was collected for an individual in a given field season. Males have names in all capitals, whereas females have names in capitals and lower case.

**Table 2.** Summary of changes observed in each chimpanzee with progressive aging. Summaries describe the differences between the first and last field season each individual was sampled (although models underlying each result used data from all field seasons). The term ‘Possible Mild Increase/Decrease’ is used to note where we identified a change for a particular metric, but this change was considerably smaller than for other individuals, and therefore could be due to chance. We address these instances on a case-by-case basis within the Results. Names of males are listed in all capitals; females’ names are in capitals and lower case.

| Individual | Sex | Attendance | Engagement | Tool Selection Time | Oil-Palm Nut Cracking |  |  | Summary |
| --- | --- | --- | --- | --- | --- | --- | --- | --- |
|  |  |  |  |  | Total Time | # Actions | # Strikes |  |
| Fana | F | Decrease | Decrease | Increase | Possible mild increase | - | - | Rate of attendance at the outdoor laboratory decreased, and Fana engaged with nuts and stones less often when present. Fana took longer to select tools. However, her general tool-using efficiency was mostly constant. |
| Jire | F | Decrease | Possible Mild Decrease | Increase | Mild increase | - | - | Rate of attendance at the outdoor laboratory decreased. Jire took longer to select stone tools. General tool-using efficiency was mostly constant, but Jire took slightly longer to crack oil-palm nuts in later years. |
| TUA | M | - | Possible Mild Increase | Decrease | - | - | - | Tua became slightly faster at selecting tools, and his tool-using efficiency was constant across years. |
| Velu | F | Decrease | Decrease (although sample size was small in 2011) | - | Increase | Increase | Increase | Rate of attendance at the outdoor laboratory decreased. Velu engaged with nuts and stones less often when present. Velu took longer to crack oil-palm nuts, and used more striking actions. |
| Yo | F | Decrease | Possible Mild Increase | Increase | Increase | Increase | Increase | Rate of attendance at the outdoor laboratory decreased. When present, Yo spent a slightly greater proportion of time engaging with nuts and stones. Yo took much longer to select tools, and demonstrated the largest reduction in tool-using efficiency. |

A summary of sampling effort for each chimpanzee in each field season can be found in Table 1, including their estimated ages in each field season (and subsequent year of death), the number of encounters sampled for behavior coding for each individual (between 10 and 1 per individual per field season; median = 10), the total duration each individual was observed during the field season (mean = 280.6 minutes per individual per field season; SD = 83.3 minutes), and the number of nuts that they were observed cracking (between 141 and 5 nuts per year; median across individuals and field seasons = 80).

### 2.2 Attendance at the outdoor laboratory

We modelled the rate at which focal old-aged chimpanzees attended the outdoor laboratory over progressive field seasons, whilst controlling for the total length of each field season. We compared this relationship with attendance data from younger individuals (between the ages of 8 and 30 years; see Fig. 2a). This younger cohort acted as a baseline control for changes in attendance rate at the population level that are unlikely to be due to senescence. In each sampled field season, the number of individuals over the age of 30 (older cohort) varied between five and six individuals, whereas the number of chimpanzees between the ages of 8 and 30 years (younger cohort) varied between five and two individuals.

**Figure 2.**
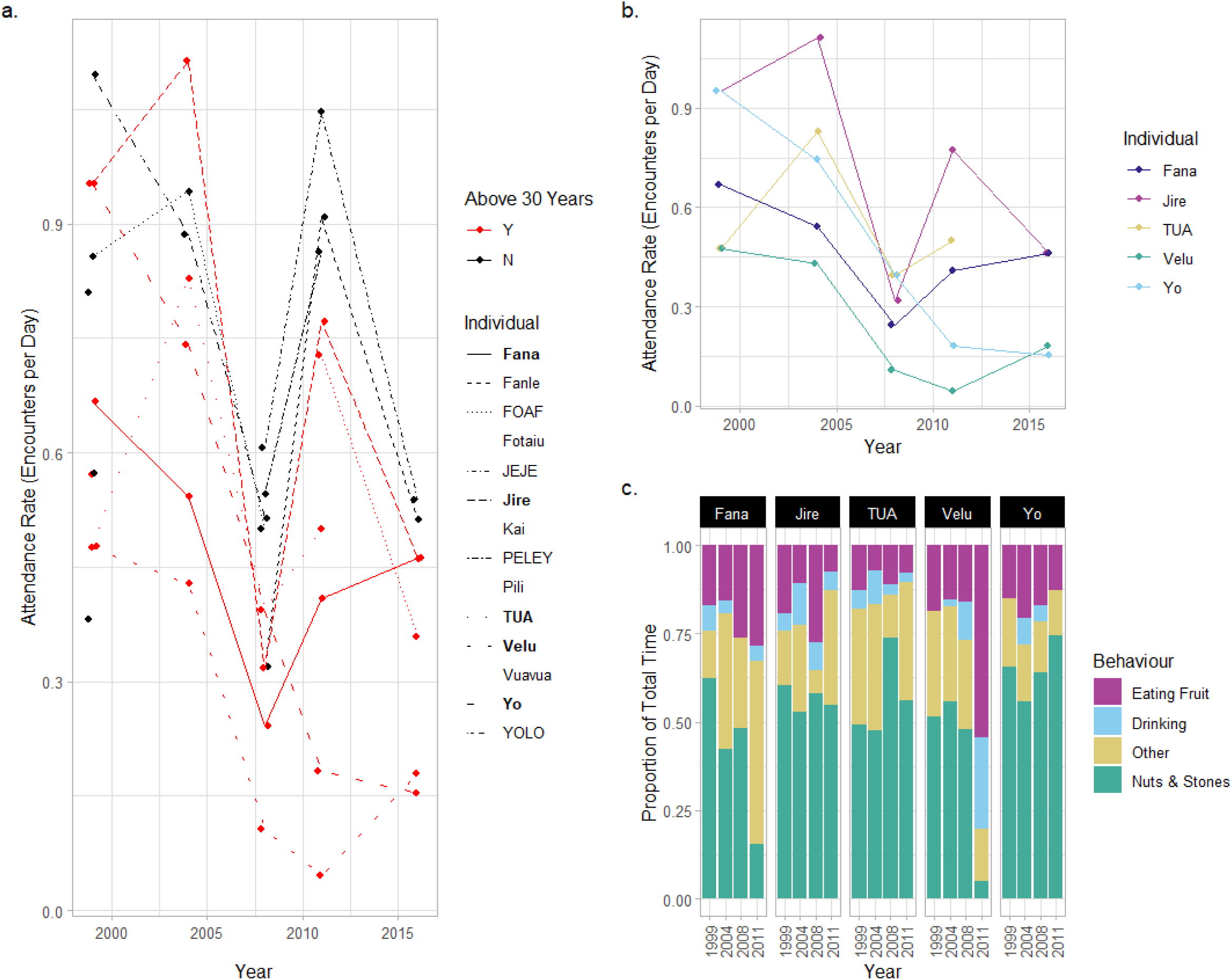
Attendance and behavior of old-aged chimpanzees at the outdoor laboratory. **(a)** Attendance rate for individuals at the outdoor laboratory between the 1999 and 2016 field seasons. Red points and lines indicate individuals are over 30 years old; black points and lines indicate individuals younger than 30 years. Lines are drawn for all individuals who attended the outdoor laboratory in two or more field seasons (individuals who were only sampled in one field season have blank spaces in the legend). The names of males are provided in all capitals, and females are provided with capital and lower-case letters. Focal old-aged individuals are indicated in the legend in bold. **(b)** Attendance data for the five focal individuals as they age from 1999 to 2016. **(c)** The proportion of total time individuals spent engaging in four different categories of behavior at the outdoor laboratory between 1999 and 2011 (data collected at the first outdoor laboratory location only).

In 1999, chimpanzees in the older cohort had a marginally lower attendance rate than younger cohort (encounters/day: older cohort = 0.68; younger cohort = 0.74). However, by 2016, this difference was much larger, with older chimpanzees being less likely to visit the outdoor laboratory compared to younger individuals (encounters/day in 2016: older cohort = 0.32; younger cohort = 0.53; see Fig. 2b for data from each elderly individual). Correspondingly, a model of attendance rate over successive field seasons (including data across all field seasons) identified a significantly greater decline in attendance rate for the older cohort across successive field seasons as compared to the younger cohort (interaction effect of field season and age-cohort: z = −2.285; p = 0.022; see Table S2 for full model output). Conversely, we found no evidence of a decline in attendance rate for chimpanzees in our younger cohort over successive field seasons (z = −1.572; p = 0.11). We confirmed the importance of this interaction effect by comparing the fit of the full model with a model lacking the interaction effect between age-cohort and field season (therefore assuming an identical relationship across field seasons for both cohorts) and a null model (which assumed no effect of field season). AIC comparison confirmed that the model which included an interaction effect offered the best explanation of the data relative to model complexity (AIC_Interaction_ = 340; AIC_Interaction-removed_ = 343; AIC_Null_ = 376). All four elderly female chimpanzees exhibited attendance rates that were lower in 2016 than in 1999; however, for Tua (the elderly adult male), attendance rates in his earliest and latest field seasons were approximately equal (1999 = 0.48 encounters/day; 2011 = 0.5 encounters/day; see Fig. 2b).

### 2.3. Behaviors at the outdoor laboratory

We measured how the proportion of time each elderly individual spent interacting with nuts and stones when present at the outdoor laboratory changed over successive field seasons (see Fig. 2c). By 2011, the final year of this analysis (we omitted the use of 2016 for this analysis, see methods), two old-aged individuals (Fana and Velu) spent substantially less time engaging with nuts and stone tools (% time engaging with nuts and stone tools in each season; Fana: 1999 = 62.3%; 2011 = 15.6%; Velu: 1999 = 51.6%; 2011 = 5.0%). We did not observe either individual successfully cracking any nuts at the outdoor laboratory in 2011. Most interactions with nuts and stone tools involved scavenging kernels from the ground (produced by other individuals’ nut cracking behaviors), or in the case of Velu, a short attempt to crack open a nut, before ceasing the behavior and feeding from oil-palm fruits. Correspondingly, in 2011, Fana spent more time engaging in ‘Other’ behaviors, such as resting and grooming (+38.3% of total time in 2011 compared with 1999), whereas Velu spent more time eating palm fruits (+35.5%) and drinking water from the water point (+25.8%; note that data for Velu come from a single encounter in 2011, so proportions should be interpreted with caution). Conversely, three individuals spent similar proportions of time engaging with nuts and stone tools when present at the outdoor laboratory (change in % time between 1999 and 2011: Jire = −5.5%; Tua = +6.9%; Yo = +8.9%). For these individuals, the small-scale changes in total time engaging with nuts and stones are more likely to be the product of chance fluctuation in engagement across field seasons.

### 2.4 Tool-selection time

Chimpanzees select stone tools based on their physical characteristics such as raw material, size, and weight^32,37,39,63^. Therefore, stone-tool selection at the central stone matrix represents a complex decision-making task where chimpanzees must evaluate an extensive set of options (over 50 total stones). Between 1999 and 2016, we recorded the duration of 108 stone-tool selection events across 49 encounters for the five focal old-aged individuals (see Fig. 3a, and Table S3). A mixed-effect model with the year of the field season as both a random intercept and random slope for each individual offered the best explanation for the total variation in the time taken to select tools (see Fig. 3b for a visualization of the random slope model). This model outperformed a null model which assumed no effect of field season (AIC_Year-RandomSlope_ = 231; AIC_Null_ = 234) as well as a model which included a fixed slope for field seasons – and therefore the same aging effect - across all individuals (where models were refit by restricted maximum likelihood to improve the accuracy of random-effect estimation; AIC_Year-Random_ = 244; AIC_Year-Fixed_ = 246; see Tables S4-S6 for model summaries). The best explanation for our data is therefore one where aging has a significant effect on tool-selection time; however, the effects of aging differ between individuals. The random-slope model identified that Yo underwent the greatest increase in stone-tool selection time across sampled field seasons, followed by Fana and then Jire. Velu exhibited relative consistency in the duration of stone-tool selection across field seasons, and the only individual to show a negative relationship between the duration of stone-tool selection and increasing years was Tua.

**Figure 3.**
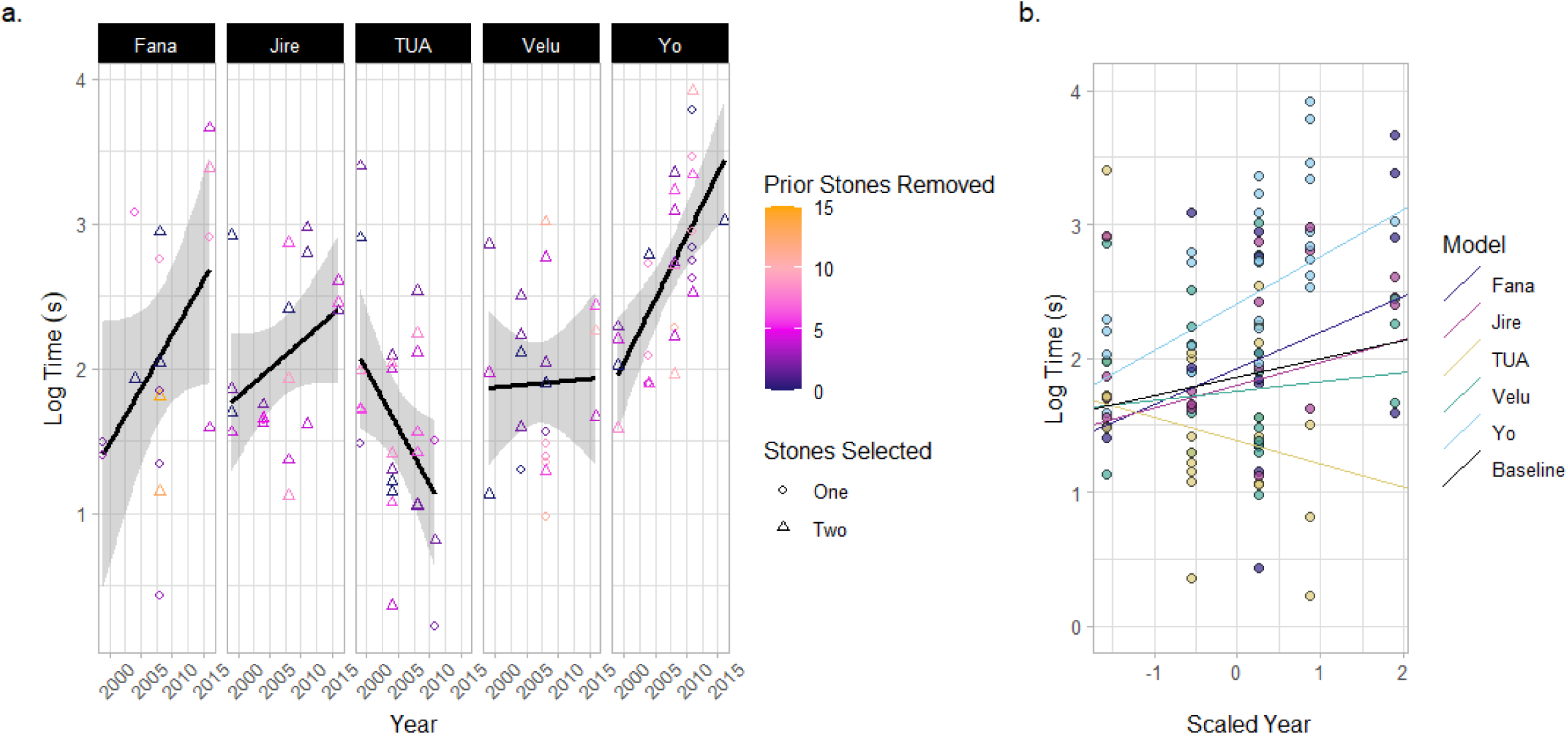
Duration of stone-tool selection events. **(a)** Tool-selection duration times for each old-aged individual. Color correlates with the number of stone tools removed from the matrix prior to that particular tool-selection event. Shape indicates the number of tools selected by an individual in a given tool-selection event. The lines and shaded areas represent a smoothed linear relationship describing the data for each individual. **(b)** A mixed effect model describing the duration of stone tool-selection events across a scaled parameter of the year for each field season. Individuals are included in the model as both a random intercept and slope. Plot shows the model’s prediction of the relationship between the duration of stone-tool selection and year for each individual, compared with the baseline fixed effect of year.

### 2.5 Efficiency processing oil-palm nuts

Across the five focal elderly chimpanzees, we recorded sequences of actions used to crack 1601 individual nuts (1538 oil-palm nuts and 63 coula nuts). As coula nuts were only provided alongside oil-palm nuts in one sampled field season (2011), we filtered data to include only the 1538 oil-palm nuts when modelling the influence of progressive old age on the efficiency of nut cracking, and describe data on the cracking of coula nuts separately (see supplementary materials).

For three efficiency metrics, models that contained the year of the field season as a fixed effect outperformed null models which assumed no change across years (models were fit using data from all field seasons). These metrics included the total time taken to crack and process nuts (AIC_Year_ = 3086, AIC_Null_ = 3092), the total number of actions used during the cracking of each nut (AIC_Year_ = 9452, AIC_Null_ = 9485), and the total number of strikes of the hammer stone performed per nut (AIC_Year_ = 6582, AIC_Null_ = 6617; see Fig. S1 for data on all measured efficiency metrics, including those where we found no effect of aging, and Tables S7-S9 for model summaries for the three metrics which exhibited changes over field seasons).

Whilst all four female chimpanzees took longer to crack nuts in later years, the extent of change varied across individuals from profound changes that are more likely to indicate senescence, to changes that were more likely attributable to small-scale chance fluctuations across nuts (see Fig. 4; see Table S10 for model coefficients for each individual). Yo exhibited the largest increase in time between 1999 and 2016 (+28.4 s; difference between 1999 and 2016 = +104%; see supplementary materials for an additional dataset for the duration of nut cracking behaviors performed by Yo in 2018, and corresponding analysis, which demonstrates that Yo took longer to crack oil-palm nuts at an even older age), and Velu exhibited the second largest increase in total time taken to crack oil-palm nuts (+26.5 s; +158%). Jire exhibited a more moderate increase in time taken to crack oil-palm nuts during the same timeframe (+9.7 s; +79%), followed by Fana, who exhibited the smallest change across all females (+5.7 s; +52%). For Fana, it is unclear whether this change represents chance fluctuations, given such small changes in nut cracking duration. Tua, the adult male, exhibited little evidence of a change in oil-palm nut cracking duration across field seasons, with his mean duration of oil-palm nut cracking decreasing by 0.67 s by 2008 (the latest year Tua was observed cracking oil-palm nuts).

**Figure 4.**
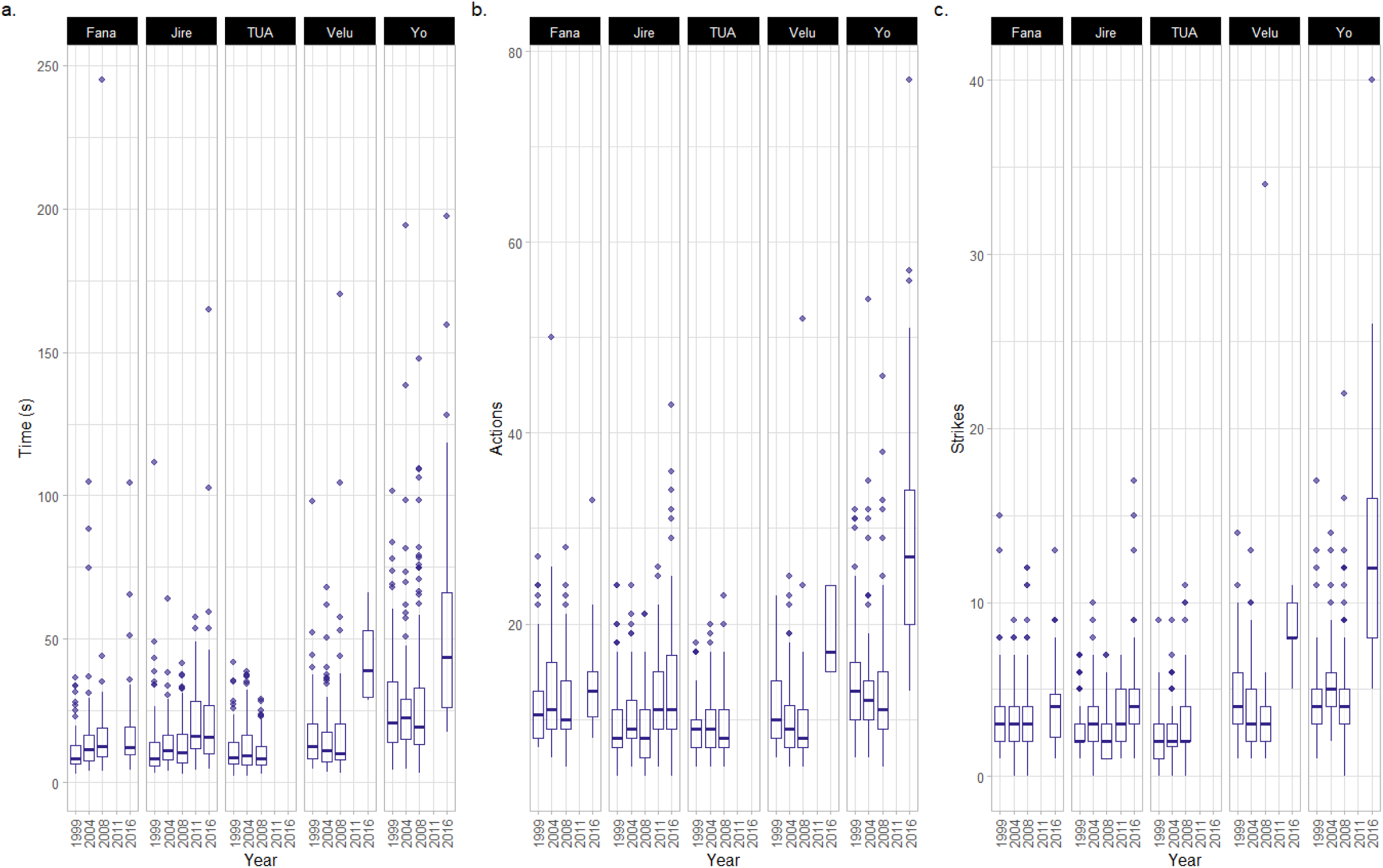
Metrics of efficiency for the cracking and processing of oil-palm nuts. This plot only includes metrics in which random slope models outperformed corresponding null models. All data relate to the cracking and processing of individual oil-palm nuts. These metrics include, for each nut cracked, (a) the total time taken (b) the number of discrete actions, and (c) the number of hammer strikes.

Between 1999 and 2016, the median number of actions Yo used to crack and process each oil-palm nut increased (+14 actions per nut; +108% change between 1999 and 2016), as did the median number of times Yo struck each nut with the hammer stone (+8 strikes per nut; +200% change between years). By analyzing the composition of Yo’s action repertoire between 1999 and 2016, we determined that, in addition to more striking actions, Yo peeled shell away from kernels using her teeth more frequently in later years (peelteeth SHELL = +6.78% of the total action repertoire composition) and also consumed all kernel in a greater number of bites (eat KERNEL = +5.16% change in total action repertoire composition). Similarly, Velu exhibited an increase in the median number of actions used to crack and process each nut between 1999 and 2016, though this effect was milder (+7 actions; +70% change between 1999 and 2016). For Velu, the majority of these additional actions were strikes of the hammer stone (+4 strikes; +100%). For Fana and Jire, we detected a smaller change in the median number of actions used to crack open oil-palm nuts by 2016 (Fana = +2.5 actions; Jire = +3 actions); however, given that these changes are small, it is more difficult to rule out the possibility that they are due to chance fluctuations in samples between years. Additionally, for Fana and Jire, the median number of striking actions performed per nut in 1999 and 2016 were very similar (Jire: +2 strikes; Fana +1 strike). We found no evidence of Tua performing more or fewer actions when cracking oil-palm nuts (including hammer strikes) over field seasons.

## 3. Discussion

We provide the first account of how a tool-use behavior performed by wild chimpanzees changed with progressively older age. Across a seventeen-year window, during which chimpanzees aged from 39-44 to 56-61 years old, elderly chimpanzees began visiting experimental nut cracking sites less frequently. This change in attendance rate was not observed for younger chimpanzees, and was therefore confined to individuals who were approaching their maximum age in the wild. In addition, when present at nut cracking sites, two individuals (Fana & Velu) exhibited a lower engagement with nuts and stone tools in 2011 (the latest field season for this analysis) as compared with their behavior in earlier field seasons. With progressive aging, three individuals took longer to select stone tools (Yo, Fana & Jire), several individuals took longer to crack open oil-palm nuts and consume all of the kernel, and two individuals (Yo and Velu) cracked and processed oil-palm nuts using a greater number of actions, including more frequent strikes of the hammer stone. Across metrics of engagement and efficiency, we detected interindividual differences in the extent to which nut cracking behaviors changed with progressive aging, including some individuals who exhibited little-to-no change in behavior across the time period sampled. Overall, our results provide initial evidence that elderly wild chimpanzees are subject to variable effects of aging on their habitual stone tool-use behaviors.

Whilst the outdoor laboratory locations are experimentally-created nut-cracking sites, previous research on the performance of nut cracking across the home range at Bossou established that chimpanzees visit the outdoor laboratory at rates comparable to naturally-occurring sites^64^, meaning that for most individuals at Bossou, the outdoor laboratory is a readily-visited site for foraging. However, at progressively older ages, elderly chimpanzees began to visit the outdoor laboratory less frequently. This result was not found for younger chimpanzees between the ages of 8 and 30 years old, suggesting that changes in elderly chimpanzees’ attendance rates were not an artefact of environmental changes over field seasons (which would likely affect both age cohorts similarly).

In addition, by 2011, two elderly chimpanzees (Velu & Fana) engaged with nuts and stone tools less frequently than they had done during visits to the outdoor laboratory throughout earlier years (though note for Velu, data for 2011 were collected from a single encounter). The possible causes for these changes in attendance rate – and engagement with nuts and stone tools when present at the outdoor laboratory - are manifold. Firstly, physiological changes with aging – such as in metabolism and nutrient requirements – could influence individuals’ dietary preferences, reducing the appeal of nuts as an available food source. Secondly, physical senescence may make visiting nut cracking sites more challenging for elderly individuals, such as through reduced capacity for locomotion. Changes in locomotion have been documented in aging captive apes^18^. In the context of wild apes, reduced mobility may further restrict the available area that elderly chimpanzees can traverse during daily ranging, encouraging them to visit foraging patches that are more proximal to their immediate locations than the outdoor laboratory. Chimpanzees habitually plan their routes towards tool-use sites^64^; however, the extent to which their doing so changes during old age is unclear. Thirdly, it is possible that tool use may become more challenging due to changes in elderly chimpanzees’ cognitive or physical condition (see latter sections of our discussion), which may reduce the appeal of visiting nut cracking sites. Fourthly, changes in social association may have influenced the likelihood that elderly chimpanzees visited experimental nut cracking sites. When experiencing increasingly old ages, many primates^18,25,29,65,66^ – and other mammals^67–69^ – display higher social selectivity, and spend a greater proportion of their time either in isolation or in smaller group associations. This social aging arguably may not be senescence *per se*, but can either be the result of senescent processes (perhaps even exacerbating senescence^67^), or may act to benefit individuals experiencing senescence, such as reducing the risk of injury from antagonistic interactions with social partners, or by shielding oneself from contagious disease^27,67^. In the context of our study, chimpanzees at Bossou typically arrive at the outdoor laboratory when travelling in groups, and therefore party members may be led to nut cracking sites by associated conspecifics, rather than through independent choice. Previous research has demonstrated that whilst gregariousness appears to be constant across ages in the Bossou chimpanzees^70^, two old-aged individuals became notably peripheral in social networks constructed using data from the 2011 field season^71^ (Yo and Velu; the two individuals we identified as having the lowest attendance rates), indicating that they were more often seen arriving at the outdoor laboratory alone or travelling as a pair. In tandem with the profound reduction in population size at Bossou throughout our study period^72^, changes in social structure may have disproportionately affected elderly chimpanzees, leading to a lower likelihood of visiting the outdoor laboratory alongside – or to regroup with – other community members. Further research is required to link our observed changes in nut cracking engagement to their underlying causes.

Through the collection of longitudinal behavioral data, we were able to evaluate whether at older ages, chimpanzees perform less streamlined tool-use behaviors. At Bossou, chimpanzees exhibit interindividual variation in nut cracking efficiency; however, within-individual efficiency is stable over the majority of each chimpanzees’ lifetime^57^. To assess for aging effects, we therefore compared behavior in middle and later adulthood for each chimpanzee, permitting signals of aging to be identified using individually-bespoke baselines of efficiency. Focal chimpanzees exhibited high efficiency in the first field season (1999) suggesting that we began sampling data before the onset of tool-use senescence (see Biro et al. 2003 for data on striking efficiency of adults at Bossou^73^, where all adults use approximately 2-4 strikes per oil-palm nut – our focal subset of adults fell within this range in 1999). Of the five individuals included in our study, three individuals took longer to select stone tools over progressing field seasons (Yo, Fana & Jire), two females showed very small increases in the amount of time taken to crack and process nuts (Fana & Jire), whereas two other females exhibited comparatively dramatic increases in the time taken to crack and process each nut (Yo & Velu). Yo and Velu also exhibited more frequent striking using the hammer stone, and Yo performed additional actions to peel shell away from kernels with her mouth, and consumed kernels with a greater number of bites (see Supplementary Video 1 for footage of Yo cracking oil-palm nuts in 1999 and 2016). Our results therefore suggest that there is measurable interindividual variation in the effects of extreme aging on the efficiency of tool selection and use in wild chimpanzees.

Our findings mirror reports of interindividual variability in the senescence of cognitive and physiological systems in both captive and wild primates^19,21,43,74,75^. Whilst it is outside of the scope of our study to determine the precise cognitive, physical, and resultant motivational changes that led to reductions in engagement with tools, rates of tool selection, and the efficiency of tool use, our results can generate predictions about how changes in fine-scale processes may translate into the behavioral changes we have observed with progressive aging. For the chimpanzees that experienced reductions in nut cracking efficiency (as discussed above), these changes could be due to changes in physical strength and dexterity^15,43^, visual acuity^23^, dentition (where tooth decay may render peeling and chewing actions more difficult^15,76^), or possibly due to changes in cognition relevant to effective tool use^10,12^, such as executive functioning or working memory. Changes in tool use behavior may also have emerged to actively compensate for the effects of senescence that are not in themselves specific to tool use; for example, tooth wear, and subsequent periodontal disease, is common in old-aged chimpanzees^15,76^. The additional strikes of the hammer stone performed by Yo and Velu to crack oil-palm nuts may have therefore been motivated by the desire to break the kernel into a greater number of smaller pieces, which were easier to peel and consume. This conclusion is somewhat supported by post-mortem data for Velu following her death in 2017, as she exhibited heavy wear patterns on both incisors and premolars, and was missing several molars on the lower jaw^77^. Similarly to tool-using efficiency, the duration of tool selection may have taken longer for several individuals due to difficulties identifying the properties of available stones, perhaps due to changes in perceptual systems (such as poorer vision^23^), or cognitive challenges in predicting the properties of objects. Alternatively, longer tool-selection times could have also been due to the need for chimpanzees to weigh up the benefits of specific tool properties (e.g. weight and size), relative to age-related changes in their physical strength and mobility.

Bridging the gap between age-related changes in tool-use behavior, and changes in cognitive and physiological processes, requires further study, and would likely benefit from experimental approaches where possible.

Our study used longitudinal video data collected from a unique, decades-long systematic study of wild chimpanzee tool use, and this video archive likely represents the only currently available data source to study intraindividual variation in chimpanzee nut cracking across the span of decades^57^. Moreover, given the exceptionally long lives of several chimpanzees at Bossou^78^, Bossou represents a unique population whose individuals can be readily studied to examine the effects of senescence in the wild at the level of the individual. However, the use of existing video data, and time-intensive methods for fine-scaled behavior coding, means that our study faced a number of limitations.

Firstly, we were unable to analyze whether specific ecological variables – such as the availability of different food sources, and demographic changes at Bossou – influence aging individuals’ attendance at the outdoor laboratory, as well as at naturally-occurring nut cracking sites (see discussion above). Nevertheless, by using the attendance of younger individuals as a control, we were able to identify that the reduction in attendance over field seasons was limited to older individuals, suggesting a specific change in behavior associated with chimpanzees experiencing progressively old age. Further research is needed to understand what factors lead to this reduced engagement with nut cracking at older ages, how aging individuals compensate for calorie loss from reduced nut cracking, and more generally, how aging influences chimpanzees’ foraging behaviors across their entire home range.

Secondly, we were not able to eliminate all possible ecological explanations for age-related changes in tool-using efficiency^79,80^. Some ecological variables were controlled for as part of the experimental set up at the outdoor laboratory. For example, chimpanzees were always tested during similar months of the year in each field season, and were provided with oil-palm nuts at a suitable stage of maturity (and thus of similar hardness). Previous studies at Bossou have also revealed that chimpanzees are able to select nuts that are suitable for cracking, further suggesting that chimpanzees can account for differences in nut quality during tool use^81^ (though this could also have been affected by aging itself, such as through reduced visual acuity impeding suitable nut selection^23^). Whilst it is possible that other ecological factors may influence our results, we believe that inter-seasonal ecological differences are unlikely to fully explain them. Firstly, if differences in chimpanzees’ nut cracking efficiency between seasons was being driven by ecological variation, we predict that this inter-annual variation would be equally present across early and late field seasons. However, contrary to this prediction, nut-cracking efficiency was very similar across early field seasons (and across individuals during these years); changes in efficiency were only detected in later field seasons, when chimpanzees were sampled at older ages. Secondly, if inter-annual differences in ecology rendered nut cracking more difficult in later field seasons, we would expect these ecological changes to affect all individuals similarly. This was also not the case, as changes in behavior differed between individuals – some maintained similar levels of efficiency (such as Fana), whereas other individuals experienced profound reductions in efficiency (such as Velu and Yo). For Yo (the individual with the greatest reduction in tool-using efficiency) the observations that Yo took even longer to crack oil-palm nuts in 2018, (and that Yo also struggled to crack coula nuts in 2011, see supplementary materials for further information) provide evidence for a directional, individual-specific reduction in nut-cracking efficiency across later years, rather than her efficiency being dictated by ecological variation. We could not provide data on a cohort of younger adults cracking nuts across field seasons, as there was only one young adult (male) individual at Bossou between 1999-2016. An additional younger adult cohort would have been a desirable control group for our study, to complement our long-term efficiency baselines we provide at the level of the individual. For future studies, we believe that the collection of a wider array of ecological data would be valuable for discriminating between explanations with greater confidence and, where possible, future studies should aim to collect long-term data on the behavior of younger adults as additional control groups.

Thirdly, as data were collected from video footage, we could not examine how the dimensions of tools and raw material types influenced chimpanzees’ tool-use behaviors over different encounters, and across field seasons. Throughout long-term data collection at Bossou, the stone tools available at the outdoor laboratory have been kept as consistent as possible, and chimpanzees select tools with physical properties which likely enhance nut cracking efficiency^32,63^. Indeed, changes in tool-using efficiency may be borne from increasing difficulty selecting appropriate tools with aging. Future studies should gather data on tool properties to support analyses of aging effects in tool use. These studies could be combined with data collection on even more fined grained behaviors of apes when using tools, such as the specific grips used to manipulate objects over time^82,83^.

Fourthly, our study was limited to a small sample size of five aging chimpanzees. This is an inevitability of collecting longitudinal data on long-lived wild individuals who are approaching the ends of their lifespan. Further data is needed – including from other field sites – to establish whether the variation in aging effects we have found generalize to other individuals and populations (and, where possible, other tool use behaviors).

Much like human technical skills, the tool-use behaviors of chimpanzees and other great apes are culturally-variable behaviors that are acquired through a mixture of social learning and individual practice^44,56,73,84^. As previously mentioned, prior research on nut cracking behavior at Bossou has revealed that chimpanzees vary in the age of skill acquisition^44,56^, and the efficiency with which nut cracking is performed across adulthood varies between individuals, yet is stable within individuals over time^57^. These differences in individual performance are, in part, likely due to differences in the environments of learners during skill acquisition^85^. Our results expand on these findings to highlight that, whilst chimpanzees at Bossou perform nut cracking behaviors throughout their lives (suggesting that this is an important skill for this population), progressive aging and senescence may exaggerate interindividual differences in the performance of socially-learnt skills, as seen in humans^86,87^. Taken as a whole, our results provide further evidence that the cultural behaviors of humans and chimpanzees exhibit marked similarities that extend across their lifetimes including, for some individuals, a period of technological senescence that can manifest across the oldest years of life. Further research should examine whether similar factors influence senescence of technical skills between humans and apes. This research should include studies that determine whether skill proficiency in early life and adulthood, and the frequency with which skills continue to be performed in later life, influence the rate and likelihood of senescence. Furthermore, debate continues as to whether human behaviors are particularly susceptible to the effects of aging due to the development of neurodegenerative diseases, such as Alzheimer’s and Parkinson’s disease^2,62,88,89^. Current evidence suggests that the effects of these neurological changes in humans have far more profound impacts on behavior, despite aging primates sometimes exhibiting similar neuropathological signatures of aging^5,90–92^. Yet, it is still not clear whether this conclusion comes from a lack of behavioral data on aging in primates. We were not able to collect neurological samples from the chimpanzees in our study, and cannot say whether the changes we observed could be associated with such processes (including for Yo, whose more profound senescence renders her the most likely to suffer from neurological decline). However, tool use could offer a suitable domain in which to examine the possible manifestation of such diseases (if present). Field sites should make specific efforts to collect long-term data on behavioral changes with aging – including for tool-use – and where possible aim to sample neurological data following the natural deaths of wild, elderly individuals. These data will provide important insights into the underlying causes of human and chimpanzee skill disruption in the contexts of healthy and diseased aging.

In foraging contexts, multi-object tool-use behaviors are often used to extract energy-rich resources that would not be otherwise accessible^93^. Given the sample size of our study, and limitations in the wider ecological data available, it was not possible to identify the influence of tool-use senescence on individual survivability through reduced energy intake. Answering this question would require data on a greater number of tool-use behaviors, spanning across the home-range. However, the two females in our study who exhibited limited changes in tool-using efficiency (Jire & Fana) had the longest lifespan after the end of our study, whereas Velu and Yo (who experienced greater difficulty cracking nuts in later years) died comparatively shortly after our sample period. While this sample is too small to make strong claims, this pattern could indicate that the senescence of tool-use behaviors, and the energy-rich foods to which they permit access, can contribute to reduced survivability in the wild. Alternatively, changes in tool-use behaviors could offer a litmus for identifying individuals experiencing profound generalized senescence in the wild, where changes in many physiological and behavioral processes subsequently impact an individual’s survivability. By extension, outside of tool use, our data on individual survivability – alongside previous studies of chimpanzees at Bossou – invites speculation surrounding whether the social relationships of specific individuals may reduce or accentuate the rate and intensity of senescence with progressive aging. In addition to living for a longer period after the final field season we sampled, Fana and Jire enjoyed more central social positions than other elderly females at Bossou^71^, and had a greater number of living relatives during the latter field seasons (during which Jire had two mature offspring, and Fana had two mature offspring, and two young grandchildren). Given that only eight individuals remained at Bossou by 2016, these familial relationships constituted a significant proportion of the overall community. Further research will provide insight into if and how lifetime sociality and familial success influences the onset and rate of senescence in wild chimpanzees, for example through socially-mediated resource acquisition.

In sum, we demonstrate that the tool-use behaviors of wild chimpanzees can change in old age, and that these changes vary between individuals. Whilst our study is observational, we argue that processes of senescence offer a promising explanation for such changes, particularly in regard to changes in individual tool-using efficiency. Our findings reflect the conclusions of previous studies investigating the cognitive and physiological senescence of primates living in captivity, and generate a myriad of further questions about the possible underlying mechanisms of tool-use senescence, as well as more general behavioral aging in wild apes. As questions surrounding aging benefit from longitudinal data – which for many primates require collection across decades – the possibility that such questions will be answered is becoming increasingly more precarious in an age where primates are at a heightened risk of extinction^94^. The conservation of wild primate populations is paramount for our understanding the dynamics and evolution of primate aging processes, which may reveal insights into the forces influencing behavioral stability and senescence across the human lifetime.

## 4. Methods

### 4.1 Study site and video archive

Established in 1976, Bossou (07° 390’ N; 008° 300’ W) is a site of long-term research focused on a small population of wild West-African chimpanzees^44^. Chimpanzees at Bossou possess an extensive repertoire of tool-use behaviors^95^, including nut cracking^32,47,55,73^. For many years, the size of the Bossou chimpanzee community remained relatively stable at around 20 individuals; however, following a flu-like epidemic in 2002 that killed 5 individuals^72^, the population began an irrecoverable decline. At the time of writing (2024), the Bossou community consists of just three individuals.

Nut cracking by Bossou chimpanzees has been studied for several decades through the use of an ‘outdoor laboratory’^47^ (established in 1988; see Fig. 1). Open during the dry season of each year (approx. November-February), the outdoor laboratory is a maintained natural clearing in the forest (approx. 7 x 20 m) where nuts and stones are provided for chimpanzees. These typically consist of oil-palm nuts, *Elaeis guineensis,* which is the only species that occurs naturally at Bossou and thus the only species of nut habitually cracked by these chimpanzees. Oil-palm nuts were sourced from local communities surrounding Bossou, and nuts provided to chimpanzees were generally of suitable maturity for cracking and consumption. In some years, coula nuts, *Coula edulis,* and in one-year panda nuts, *Panda oleosa*, were also provided. Neither coula nor panda nuts occur naturally at Bossou, but both are cracked by chimpanzees at nearby sites^73^. In addition to nuts, chimpanzees are also provided with over 50 stones organized in a square matrix in the center of the clearing at the start of each session (see Fig. 1d), as well as a raceme of edible oil-palm fruits (see Fig. 1c). The outdoor laboratory also features a water point (a hollowed section of a tree trunk; see Fig. 1b) from which chimpanzees may extract water for drinking using leaf tools. Chimpanzees visit the outdoor laboratory spontaneously as part of their daily ranging behaviors, and with comparable frequency to other naturally-occurring nut cracking sites within the home range at Bossou^64^. Upon arrival at the outdoor laboratory, chimpanzees’ behaviors are recorded by observers using 2 or 3 video cameras located behind a screen of vegetation.

Video data collected at the outdoor laboratory has recently been collated into a longitudinal archive spanning from 1990 to the present day^57,71,96^. Given the long-term use of the outdoor laboratory to study nut cracking, all chimpanzees were well habituated to the experimental set up by 1999 (the first field season from which data was sampled for this study). The Bossou archive also contains data on the composition of all recorded encounters with groups of chimpanzees at the outdoor laboratory across this timeframe, and can be combined with recorded family lineage data (including birth data for individuals born after the establishment of long-term research at Bossou). The Bossou archive therefore presents a novel opportunity for longitudinal analysis of whether and how tool-use behaviors change across the later periods of chimpanzee lifespans.

We collected and analyzed data on the behaviors of chimpanzees at the outdoor laboratory at Bossou, spanning a seventeen-year period (1999-2016). This 17-year period contains the majority of the available video data collected at Bossou. To balance fine-grained behavior coding (which is highly time intensive) with the need to sample data over an extended time period to capture possible effects of long-term senescence, we aimed to sample field seasons within this time window at four-year intervals; however, in some years, data was not collected at Bossou (such as during years where disease outbreaks were affecting local communities). In such instances, we sampled the closest field season which likely contained sufficient data for our analyses. Thus, during this period, we sampled data from five field seasons (1999, 2004, 2008, 2011, and 2016).

Over the years, the outdoor laboratory has operated at two locations in the forest. Between 1999 and 2008, data was only collected at one specific location across the years. However, in 2011 a second clearing was made available to chimpanzees, with an identical experimental set up. In 2011, data was collected in both locations simultaneously. By 2016, all data collection had transitioned to the newer, second location. For the majority of our analyses, we therefore used data from the first location between the years of 1999-2011, and the second location in 2016. For only one analysis (attendance rate, see below) we used data from both the first and second outdoor laboratory location in 2011. All behavior coding in our study was performed using the open-access software BORIS^97^, with videos viewed on a 59cm monitor.

### 4.2 Attendance

We recorded the total number of times chimpanzees were encountered at the outdoor laboratory in each field season. An encounter was defined as any instance where one or a group of chimpanzees visited the outdoor laboratory, and lasted until all chimpanzees left. If a chimpanzee left the outdoor laboratory and returned before all other individuals had left, it was classed as part of the same encounter. We collected data on the five chimpanzees experiencing increasingly old age, as well as for an additional cohort of all adult and subadult individuals between the ages of 8 and 30 years old. This second cohort of individuals allowed us to evaluate whether changes in attendance rate over progressive field seasons may be due to confounding factors other than senescence, such as changes in environmental food availability or population structure, or changes in the number of experimental nut cracking sites. For our younger cohort of chimpanzees, we used a minimum age of 8 years old, as chimpanzees at Bossou acquire the skill of nut cracking by 7 years of age. For this analysis, we also excluded two adult females (Pama and Nina) who never learnt to successfully crack nuts using stone tools. To evaluate whether progressive aging reduced attendance rates, we constructed a Poisson GLMM with field season as a continuous fixed effect (scaled about the mean), and cohort (above or below 30 years) as a categorical variable. Rates of attendance were estimated using encounters (a count variable) and the inclusion of an offset term in our model (the number of days of the field season). Importantly, we included an interaction term in our model to see whether the relationship between aging across field seasons and attendance rate was more severe for our focal elderly individuals compared with the younger cohort. Individual ID was included as a random intercept to account for repeated measures of the same individual across field seasons.

### 4.3 Behaviors at the outdoor laboratory

We estimated whether, over successive field seasons, elderly individuals began spending a greater or lesser proportion of their time engaging in nut cracking when present at the outdoor laboratory, as compared with other commonly performed behaviors. For each elderly individual, we sampled the first 10 encounters of each field season in which they were present (totaling a maximum of 50 encounters per individual across all field seasons). For each encounter, we timed how long individuals engaged in one of four mutually exclusive categories of behavior:

1. **Engaging with Nuts and Stone Tools**: the total amount of time individuals spent interacting with nuts, nut fragments (including shells and kernels), and stones. Coding began when an individual first touched a nut, nut fragment, or stone.
2. **Drinking Water**: the total amount of time individuals spent producing leaf tools and drinking from the water point. Coding began when an individual stripped leaves from branches to begin manufacturing a leaf tool to aid in drinking water, or, if reusing a tool that had previously been made and discarded, when they collected the tool, and began to approach the water point to drink.
3. **Eating Oil-Palm Fruits**: the total amount of time individuals spent consuming oil-palm fruits from the raceme. Coding began whenever an individual started picking oil-palm fruits from the raceme, or collecting and consuming oil-palm fruits off of the ground.
4. **Other**: any other behaviors (e.g. grooming, playing, resting).

For behaviors 1-3, coding ceased when individuals stopped interacting with all relevant objects of one behavior (e.g. leaf tools, stones, fruits), and began engaging in another behavior (either another behavior from 1-3, or ‘Other’ behaviors such as grooming). If an individual dropped all relevant items and was idle for at least one minute, they were considered to be resting, and behaviors were marked as ‘Other’. For field seasons in which individuals did not visit the outdoor laboratory on at least 10 occasions, all available encounters were used. We also recorded the total amount of time each focal individual spent at the outdoor laboratory during each encounter. We calculated the proportion of time individuals spent engaging in each type of behavior across all encounters of each field season (thus, controlling for differences in the total time individuals were observed between years). For this analysis, we omitted any videos collected at the second outdoor laboratory site (including some videos from 2011 and all videos from 2016). We chose to do so as the two sites vary in how accessible they are (with the original site being situated at the top of a steep hill, but the second requiring no climb). This difference in terrain may influence the behaviors of the chimpanzees once arriving at the outdoor laboratory, for example, more time resting following climbing.

### 4.4 Stone-tool selection

We measured the time taken for the old-aged chimpanzees to select stone tools at the central matrix during each of the sampled field seasons. We sampled the first 10 encounters with each individual for each field season. All tool-selection events in these encounters were sampled. We began timing when chimpanzees approached the matrix and we considered their gaze to be fixed on at least one stone tool. We stopped timing when chimpanzees turned away from the matrix holding all acquired stone tools. We recorded the number of stone tools taken by the focal chimpanzee during each instance of stone-tool selection (a categorical score of ‘one’ or ‘two’ stones). Additionally, we recorded the number of stone tools that had been removed from the stone-tool matrix prior to each stone selection event (including those removed by other individuals, as well as the focal chimpanzee during previous instances of stone-tool selection). We could therefore incorporate into our analysis a metric of how many options were available to chimpanzees during each stone-tool selection event (where higher numbers of previously selected tools indicated a smaller pool of possible choices). We did not sample instances where chimpanzees selected and used tools which were not positioned at the central matrix (i.e. when using tools which had already been transported away from the matrix by other chimpanzees). When modelling tool-selection time over field seasons, both the number of stones that a chimpanzee selected from the central matrix (one or two) and the number of stones previously removed from the matrix were included in our model as fixed effects, to control for additional variation in selection time introduced by these two factors (see section 4.6 below for more details on GLMM model structures).

### 4.5 Coding discrete actions

When performing complex technical skills—including tool-use behaviors— experienced apes organize individual actions into more streamlined goal-directed sequences^30,33,56,98^, and begin to successfully perform behaviors more quickly^33,37,73,82^; however, whether experiencing progressively old age influences this efficiency has not yet been studied.

To evaluate the efficiency with which focal chimpanzees engaged in nut cracking, we coded the individual, fine-grained actions used by focal chimpanzees when cracking nuts at the outdoor laboratories. We sampled videos chronologically from each sampled field season. Fine-grained action coding was conducted for each individual in each field season until at least 1000 actions had been collected, and included sequences of actions that described the cracking of at least 20 nuts in their entirety. To code sequences of actions, we used an ethogram of 34 manipulations (e.g. ‘Grasp’, ‘Drop’, ‘Strike’) and 6 objects (see Table S1 for full ethogram^33^; all discrete-action coding was performed by one individual: EHS; our ethogram was tested to ensure that it can be applied objectively as part of a previous research project^33^). The six possible objects included: Hammer, Anvil, Nut, Kernel (the edible interior of a nut), Shell (the inedible exterior of a nut), and Bare Hand (for use when individuals performed an action in the absence of an object, e.g., striking the anvil with bare hand). Stone tools were categorized as ‘Hammer’ or ‘Anvil’ based on how they were used by the individual. If an individual swapped over their hammer and anvil stones, the stones were reassigned accordingly. Thus, stone tools were coded according to their function at any specific point in a sequence. We considered an individual action to be the combination of both the manipulation performed by the chimpanzee, and the object being manipulated (e.g. ‘Grasp Hammer’ = one action).

Coding of action sequences started when an individual began interacting with a nut, nut fragment, or stone, and ceased when an individual dropped all relevant objects and either began engaging in an alternative behavior (e.g. grooming, feeding on oil-palm fruits, etc.) or rested for longer than one minute. Actions were coded as discrete events during behavioral observation, and the time of each action was automatically recorded in BORIS.

The total corpus of actions was parsed into sequences directed at the cracking of individual nuts and the consumption of all associated kernel. Only sequences which described the entire processing of a nut, from acquisition of the nut to complete consumption of the kernel, were included in our analysis. A number of metrics of efficiency were extracted from action sequences directed at individual nuts, which emphasize the speed with which kernels can be acquired and consumed, and the extent to which the actions used to do so reflect a streamlined behavioral sequence:

1. **Total time to crack nuts and consume all kernel.** The total time between the start of the first action and the end of the last action performed on a specific nut, including all actions involving the consumption of kernel, and disposal of shell fragments.
2. **Number of actions used to process nuts and consume all kernel.** The total number of actions performed during the cracking of a specific nut, including consumption of kernel and disposal of shell fragments.
3. **Number of unique types of action used to process nuts and consume all kernel.** The number of different types of actions performed by an individual when cracking a specific nut, including all kernel consumption and disposal of shell fragments. This metric is different to [2] as it describes the variety of actions employed, rather than the number; e.g. for the action sequence *A,A,B,C,B,C*, where letters denote hypothetical actions, the number of actions is six, and the number of unique action types is three {*A*, *B*, *C*}.
4. **Number of times the nut was (re)placed on the anvil**. The number of times the nut was placed on the anvil during initial placement, and also all replacements when the nut rolled off the anvil. An understanding of how to orient the anvil, and where to place the nut can reduce the likelihood of needing to perform nut replacements. Therefore, fewer nut placements reflects a greater efficiency in nut cracking.
5. **Number of strikes of the hammer stone per nut.** The number of individual strikes of the hammer stone performed on each nut that a chimpanzee cracked. This is a common metric for efficiency in chimpanzee nut cracking^73,99^, where fewer strikes indicate higher efficiency.
6. **Number of tool reorientations per nut.** The total number of horizontal rotations (‘reorient’) and vertical rotations (‘flip’) of hammer and anvil stones. Chimpanzees occasionally reorient tools during tool use, with these adjustments likely reflecting attempts to achieve greater efficiency. Once a suitable configuration of tool objects is acquired, such reorientations are usually less common.
7. **Tool changes per nut.** The number of times an individual swapped over their hammer and anvil stones, swapped out at least one stone tool for a new stone, or moved across the outdoor laboratory to use a hammer-anvil set previously abandoned by another user.

### 4.6 Statistical analyses

Statistical analyses were performed in R (version 4.3.3 Angel Food Cake). For attendance rates, tool-selection times, and the metrics of efficiency cracking individual nuts, we used linear and generalized linear mixed-effect models to assess for the effects of age (using the lme4 package^100^). Linear mixed effect models were used in any instance where time was a response variable (time was logged to confer normality); whereas, if a response variable was measured in counts (e.g., number of actions used to crack open a nut; number of encounters over a field season, etc.), we employed generalized linear mixed-effect models using a Poisson distribution. In both instances, year of the field season (scaled about the mean) was used as a fixed effect, as a proxy for age.

Our model for attendance data is described above in section 4.2 (we describe this model separately as we modelled the effect of aging in two separate groups – young and old – rather than for specific individuals). For all other metrics, we modelled the effect of aging on focal, elderly chimpanzees by using random intercepts and slopes to evaluate the effect of the increasing year of each field season. Our decision to include a random slope is supported by previous research on captive and wild primates, which suggests that the effects of senescence are highly variable across individuals for a number of physiological and cognitive processes^19,21,43,74,75^. Across metrics of engagement and efficiency, we compared random-slope models with fixed slope models - which assume that the effect of increasing field season will be the same for all individuals - as well as with null models which fit an intercept across all field seasons (and therefore assumed no effect of aging). This allowed us to evaluate whether the effect of increasing year of each field season influenced chimpanzee behavior, and also whether this effect was variable across individuals. All model comparisons were performed using AIC, where lower scores indicate better fit relative to model complexity. Where necessary, we also included the specific encounter as an additional random intercept (fixed slope) to avoid pseudoreplication from multiple tool-selection events within the same encounter.

When analyzing metrics of nut cracking efficiency (for which we had a total of seven metrics, see above), we restricted the use of models to describe changes in metrics based on whether, for at least one individual, the median value for a particular field season fell outside the interquartile range of any previous field season. If a model identified that an efficiency metric was changing across field seasons, we used model coefficients to gauge the direction of change for each individual. To estimate the magnitude of change across field seasons, we compared data collected for each individual in the earliest and latest year that they were sampled. We calculated % change as the change in a metric between the last and first field season, divided by the value for the first field season (to scale the change), multiplied by 100 (see associated code).

### 4.7 Ethics Statement

Data collection at Bossou has been conducted by different researchers over several decades. Research authorization and ethical approval were given to T.M. (who oversaw data collection during the study period covered by this project) through a memorandum of understanding between Kyoto University, and the Guinean authorities (Direction Générale de la Recherche Scientifique et de l’Innovation Technologique, DGERSIT, and the Institut de Recherche Environnementale de Bossou, IREB), to which all researchers conformed. The present study was entirely conducted using existing video data from Bossou (the Bossou Archive), and therefore did not involve collection of additional data from wild chimpanzees.

## Acknowledgments

We thank Daniel Schofield for their help in curating the Bossou video archive, and Alex Mielke for their statistical advice. For both assistance and permission to conduct research at Bossou, we thank the Direction Générale de la Recherche Scientifique et de l’Innovation Technologique (DGERSIT) and the Institut de Recherche Environnementale de Bossou (IREB), République de Guinée. For their invaluable help in the field, we would also like to thank the research assistants Boniface Zogbila, Gouanou Zogbila, Tino Zogbila, Henry Didier Camara, Marcel Doré, Jules Doré, Pascal Goumy, Gouanou Goumy, Paquile Cherif, Cé Vincent Mamy, Dagouka Samy as well as all other guides. Additionally, we thank all KUPRI-International researchers (especially Misato Hayashi, Claudia Sousa, Kimberly Hockings, Katelijne Koops, Tatiana Humle, Gaku Ohashi, Hiroyuki Takemoto, Gen Yamakoshi, Yukimaru Sugiyama, and Jérémy Koman), and IREB researchers (especially Mamadou Diakité, Makan Kourouma, and Aly-Gaspard Soumah) who contributed to the wider program of data collection at Bossou during the period of our study.

## References

1. Colman, R. J. Non-human primates as a model for aging. BBA-Mol. Basis. Dis. 1864, 2733–2741 (2018).

2. Edler, M. K., Munger, E. L., Groetz, H. & Raghanti, M. A. Nonhuman primates as models for aging and Alzheimer’s disease. in Assessments, Treatments and Modeling in Aging and Neurological Disease (eds. Martin, C. R., Preedy, V. R. & Rajendram, R.) 527–537 (Elsevier, 2021). doi:10.1016/B978-0-12-818000-6.00047-0.

3. Gilleard, C. Aging in Non-Human Primate Society: What Relevance for Social Gerontology? A&A 44, 37–56 (2023).

4. Bizon, J. L. & Woods, A. Animal Models of Human Cognitive Aging. (Humana Press, Totowa, NJ, 2009).

5. Frye, B. M. et al. Early Alzheimer’s disease-like reductions in gray matter and cognitive function with aging in nonhuman primates. A&D Transl. Res. & Clin. Interv. 8, (2022).

6. Lane, M. Nonhuman primate models in biogerontology. Exp. Gerontol. 35, 533–541 (2000).

7. Comrie, A. E., Gray, D. T., Smith, A. C. & Barnes, C. A. Different macaque models of cognitive aging exhibit task-dependent behavioral disparities. Behav. Brain Res. 344, 110–119 (2018).

8. Gray, D. T. & Barnes, C. A. Experiments in macaque monkeys provide critical insights into age-associated changes in cognitive and sensory function. Proc. Natl. Acad. Sci. U.S.A. 116, 26247– 26254 (2019).

9. Herndon, J. G. & Lacreuse, A. The Rhesus Monkey Model as a Heuristic Resource in Cognitive Aging Research. in Aging in Nonhuman Primates (eds. Erwin, J. M. & Hof, P. R.) 178–195 (Karger, 2002).

10. Lacreuse, A., Raz, N., Schmidtke, D., Hopkins, W. D. & Herndon, J. G. Age-related decline in executive function as a hallmark of cognitive ageing in primates: an overview of cognitive and neurobiological studies. Phil. Trans. R. Soc. B 375, 20190618 (2020).

11. Lacreuse, A., Parr, L., Chennareddi, L. & Herndon, J. G. Age-related decline in cognitive flexibility in female chimpanzees. Neurobiol. Aging 72, 83–88 (2018).

12. Lacreuse, A., Russell, J. L., Hopkins, W. D. & Herndon, J. G. Cognitive and motor aging in female chimpanzees. Neurobiol. Aging 35, 623–632 (2014).

13. Colman, R. & Binkley, N. Skeletal Aging in Macaque Monkeys. in Aging in Nonhuman Primates (eds. Erwin, J. M. & Hof, P. R.) 32–37 (Karger, 2002).

14. Didier, E. S. et al. Contributions of Nonhuman Primates to Research on Aging. Vet. Pathol. 53, 277–290 (2016).

15. Lowenstine, L. J., McManamon, R. & Terio, K. A. Comparative Pathology of Aging Great Apes: Bonobos, Chimpanzees, Gorillas, and Orangutans. Vet. Pathol. 53, 250–276 (2016).

16. Havercamp, K., Morimura, N. & Hirata, S. Sleep Patterns of Aging Chimpanzees (*Pan troglodytes*). Int. J. Primatol. 42, 89–104 (2021).

17. Hozer, C., Pifferi, F., Aujard, F. & Perret, M. The Biological Clock in Gray Mouse Lemur: Adaptive, Evolutionary and Aging Considerations in an Emerging Non-human Primate Model. Front. Physiol. 10, 1033 (2019).

18. Neal Webb, S. J., Hau, J., Lambeth, S. P. & Schapiro, S. J. Differences in Behavior Between Elderly and Nonelderly Captive Chimpanzees and the Effects of the Social Environment. J. Am. Assoc. Lab. Anim. Sci. 58, 783–789 (2019).

19. Tarou, L. R., Bloomsmith, M., Hoff, M., Erwin, J. & Maple, T. The Behavior of Aged Great Apes. In Aging in Nonhuman Primates (eds. Erwin, J. M. & Hof, P. R.) 209–231 (Karger, 2002).

20. Zhdanova, I. V. et al. Aging of Intrinsic Circadian Rhythms and Sleep in a Diurnal Nonhuman Primate, *Macaca mulatta*. J. Biol. Rhythms 26, 149–159 (2011).

21. Rothwell, E. S. et al. The marmoset as an important primate model for longitudinal studies of neurocognitive aging. Am. J. Primatol. 83, (2021).

22. Tardif, S. D. & Ross, C. N. Aging in nonhuman primates. in Handbook of the Biology of Aging (eds. Musi, N. & Hornsby, P.) 237–248 (Elsevier, 2021). doi:10.1016/B978-0-12-815962-0.00011-1.

23. Fujisawa, M. et al. Farsightedness (presbyopia) in a wild elderly chimpanzee: The first report. Geriatr. Gerontol. Int. 10, 113–114 (2010).

24. Newman, L. E. et al. The biology of aging in a social world: Insights from free-ranging rhesus macaques. Neurosci. Biobehav. R. 154, 105424 (2023).

25. Siracusa, E. R. et al. Within-individual changes reveal increasing social selectivity with age in rhesus macaques. Proc. Natl. Acad. Sci. U.S.A. 119, e2209180119 (2022).

26. Thompson, M. E. et al. Evaluating the impact of physical frailty during ageing in wild chimpanzees (*Pan troglodytes schweinfurthii*). Phil. Trans. R. Soc. B 375, 20190607 (2020).

27. Siracusa, E. R. et al. Social ageing can protect against infectious disease in a group-living primate. Phil. Trans. R. Soc. B 379, 20220462 (2024).

28. Cole, M. F. et al. Age-related physiological dysregulation progresses slowly in semi-free-ranging chimpanzees. Evol. Med. Pub. Health eoae010 (2024) doi:10.1093/emph/eoae010.

29. Campos, F. A. et al. Wild capuchin monkeys as a model system for investigating the social and ecological determinants of ageing. Phil. Trans. R. Soc. B 379, 20230482 (2024).

30. Byrne, R. W., Sanz, C. M. & Morgan, D. B. Chimpanzees plan their tool use. in Tool Use in Animals (eds. Sanz, C. M., Call, J. & Boesch, C.) 48–64 (Cambridge University Press, 2013). doi:10.1017/CBO9780511894800.004.

31. Call, J. Three ingredients for becoming a creative tool user. in Tool Use in Animals (eds. Sanz, C. M., Call, J. & Boesch, C.) 3–20 (Cambridge University Press, 2013). doi:10.1017/CBO9780511894800.002.

32. Carvalho, S., Cunha, E., Sousa, C. & Matsuzawa, T. Chaînes opératoires and resource-exploitation strategies in chimpanzee (*Pan troglodytes*) nut cracking. J. Hum. Evol. 55, 148–163 (2008).

33. Howard-Spink, E. et al. Nonadjacent dependencies and sequential structure of chimpanzee action during a natural tool-use task. PeerJ 12, e18484 (2024).

34. Hayashi, M. Perspectives on object manipulation and action grammar for percussive actions in primates. Philosophical Transactions of the Royal Society B: Biological Sciences 370, (2015).

35. Matsuzawa, T. Chimpanzee intelligence in nature and in captivity: Isomorphism of symbol use and tool use. in Great Ape Societies (eds. McGrew, W. C., Marchant, L. F. & Nishida, T.) 196– 209 (Cambridge University Press, Cambridge, UK, 1996).

36. Sanz, C. M. & Morgan, D. B. The Complexity of Chimpanzee Tool-Use Behaviors. in The mind of the chimpanzee: Ecological and experimental perspectives (eds. Lonsdorf, E. V., Ross, S. R. & Matsuzawa, T.) 127–140 (The University of Chicago Press, Chicago, US, 2010).

37. Boesch, C. & Boesch, H. Optimisation of Nut-Cracking With Natural Hammers By Wild Chimpanzees. Behav. 83, 265–286 (1983).

38. Bril, B., Dietrich, G., Foucart, J., Fuwa, K. & Hirata, S. Tool use as a way to assess cognition: How do captive chimpanzees handle the weight of the hammer when cracking a nut? Anim. Cogn. 12, 217–235 (2009).

39. Sirianni, G., Mundry, R. & Boesch, C. When to choose which tool: multidimensional and conditional selection of nut-cracking hammers in wild chimpanzees. Anim. Behav. 100, 152–165 (2015).

40. Glavis-Bloom, C., Vanderlip, C. R. & Reynolds, J. H. Age-Related Learning and Working Memory Impairment in the Common Marmoset. J. Neurosci. 42, 8870–8880 (2022).

41. Nichols, K. & Zihlman, A. Skeletal and Dental Evidence of Aging in Captive Western Lowland Gorillas: A Preliminary Report. in Aging in Nonhuman Primates (eds. Erwin, J. M. & Hof, P. R.) 22–31 (Karger, 2002).

42. Cann, J. A. Nonhuman Primate Models of Human Disease. in The Nonhuman Primate in Nonclinical Drug Development and Safety Assessment (eds. Bluemel, J. & Schenck, E.) 257–277 (Elsevier, 2015). doi:10.1016/B978-0-12-417144-2.00013-5.

43. Morbeck, M., Galloway, A. & Sumner, D. Getting Old at Gombe: Skeletal Aging in Wild-Ranging Chimpanzees. in Aging in Nonhuman Primates (eds. Erwin, J. M. & Hof, P. R.) 48–62 (Karger, 2002).

44. Matsuzawa, T., Humle, T. & Sugiyama, Y. The Chimpanzees of Bossou and Nimba. (Springer Tokyo, Tokyo, Japan, 2011).

45. Sugiyama, Y. Observations on the population dynamics and behavior of wild chimpanzees at Bossou, Guinea, in 1979-1980. Primates 22, 435–444 (1981).

46. Carvalho, S. & McGrew, W. The origins of the Oldowan: Why chimpanzees (Pan troglodytes) still are good models for technological evolution in Africa. in Stone Tools and Fossil Bones (ed. Domínguez-Rodrigo, M.) 201–221 (Cambridge University Press, 2012).

47. Matsuzawa, T. Field Experiments on use of Stone Tools in the wild. in Chimpanzee Cultures (eds. Wrangham, R. W., McGrew, W. C., De Waal, F. B. M. & Heltne, P. G.) 351–370 (Harvard University Press, 1994).

48. Falótico, T., Proffitt, T., Ottoni, E. B., Staff, R. A. & Haslam, M. Three thousand years of wild capuchin stone tool use. *Nat*. Ecol. Evol. 3, 1034–1038 (2019).

49. Fragaszy, D. M. et al. The development of expertise at cracking palm nuts by wild bearded capuchin monkeys, Sapajus libidinosus. Anim. Behav. 197, 1–14 (2023).

50. Falótico, T., Macedo, A. C., De Jesus, M. A., Espinola, T. & Valença, T. Nut-cracking success and efficiency in two wild capuchin monkey populations. R. Soc. Open Sci. 11, 240161 (2024).

51. Goldsborough, Z., Crofoot, M. C. & Barrett, B. J. Male-biased stone tool use by wild white-faced capuchins (*Cebus capucinus imitator*). Am. J. Primatol. 86, e23594 (2024).

52. Goldsborough, Z., Carlson, M., Reetz, L., Crofoot, Margaret. C. & Barrett, B. J. Development and social dynamics of stone tool use in white-faced capuchin monkeys. Preprint at 10.1101/2025.04.08.647785 (2025).

53. Luncz, L. V. et al. Group-specific archaeological signatures of stone tool use in wild macaques. eLife 8, e46961 (2019).

54. Proffitt, T. et al. Analysis of wild macaque stone tools used to crack oil palm nuts. R. Soc. Open Sci. 5, 171904 (2018).

55. Biro, D., Sousa, C. & Matsuzawa, T. Ontogeny and cultural propagation of tool use by wild chimpanzees at Bossou, Guinea: Case studies in nut cracking and leaf folding. in Cognitive Development in Chimpanzees (eds. Matsuzawa, T., Tomonaga, M. & Masayuki, T.) 476–508 (Springer Tokyo, Tokyo, Japan, 2006).

56. Inoue-Nakamura, N. & Matsuzawa, T. Development of Stone Tool Use by Wild Chimpanzees (*Pan troglodytes*). J. Comp. Psychol. 111, 159–173 (1997).

57. Berdugo, S., Cohen, E., Davis, A. J., Matsuzawa, T. & Carvalho, S. Reliable long-term individual variation in wild chimpanzee technological efficiency. *Nat*. Hum. Behav. (2024) doi:10.1038/s41562-024-02071-8.

58. Matsuzawa, T. Nesting cups and metatools in chimpanzees. Behav. Brain Sci. 14, 570–571 (1991).

59. Wood, B. M. et al. Demographic and hormonal evidence for menopause in wild chimpanzees. Science 382, eadd5473 (2023).

60. Sousa, C., Biro, D. & Matsuzawa, T. Leaf-tool use for drinking water by wild chimpanzees (*Pan troglodytes*): Acquisition patterns and handedness. Anim. Cogn. 12, (2009).

61. Campos, F. A. et al. Female reproductive aging in seven primate species: Patterns and consequences. Proc. Natl. Acad. Sci. U.S.A. 119, e2117669119 (2022).

62. Finch, C. E. & Austad, S. N. Commentary: is Alzheimer’s disease uniquely human? Neurobiol. Aging 36, 553–555 (2015).

63. Braun, D. R. et al. Stone selection by wild chimpanzees shares patterns with Oldowan hominins. J. Hum. Evol. 199, 103625 (2025).

64. Almeida-Warren, K., Camara, H. D., Matsuzawa, T. & Carvalho, S. Landscaping the Behavioural Ecology of Primate Stone Tool Use. Int. J. Primatol. 43, 885–912 (2022).

65. Almeling, L., Hammerschmidt, K., Sennhenn-Reulen, H., Freund, A. M. & Fischer, J. Motivational Shifts in Aging Monkeys and the Origins of Social Selectivity. Curr. Biol. 26, 1744–1749 (2016).

66. Machanda, Z. P. & Rosati, A. G. Shifting sociality during primate ageing. Phil. Trans. R. Soc. B 375, 20190620 (2020).

67. Siracusa, E. R., Higham, J. P., Snyder-Mackler, N. & Brent, L. J. N. Social ageing: exploring the drivers of late-life changes in social behaviour in mammals. Biol. Lett. 18, 20210643. (2022).

68. Rudd, L. F., Packer, C., Biro, D., Firth, J. A. & Albery, G. F. Sex-specific social aging in wild African lions. Curr. Biol. S0960982224009424 (2024) doi:10.1016/j.cub.2024.07.040.

69. Albery, G. F. et al. Ageing red deer alter their spatial behaviour and become less social. *Nat*. Ecol. Evol. 6, 1231–1238 (2022).

70. Schofield, D. P. et al. Automated face recognition using deep neural networks produces robust primate social networks and sociality measures. Methods Ecol. Evol. 14, 1937–1951 (2023).

71. Schofield, D. et al. Chimpanzee face recognition from videos in the wild using deep learning. Sci. Adv. 5, 1–10 (2019).

72. Matsuzawa, T. et al. Wild chimpanzees at Bossou-Nimba: Deaths through a flu-like epidemic in 2003 and the green corridor project. Primate Res. 20, 45–55 (2004).

73. Biro, D. et al. Cultural innovation and transmission of tool use in wild chimpanzees: Evidence from field experiments. Anim. Cogn. 6, 213–223 (2003).

74. Freire-Cobo, C., et al. Comparative neuropathology in aging primates: A perspective. Am. J. Primatol. 83, (2021).

75. Hämäläinen, A., Heistermann, M. & Kraus, C. The stress of growing old: sex- and season-specific effects of age on allostatic load in wild grey mouse lemurs. Oecologia 178, 1063–1075 (2015).

76. Albrecht, A., Behringer, V., Zierau, O. & Hannig, C. Dental findings in wild great apes from macerated skull analysis. Am. J. Primatol. 86, e23581 (2024).

77. Matsuzawa, T. Chimpanzee Velu: the wild chimpanzee who passed away at the estimated age of 58. Primates 59, 107–111 (2018).

78. Emery Thompson, M., et al. Aging and Fertility Patterns in Wild Chimpanzees Provide Insights into the Evolution of Menopause. Curr. Biol. 17, 2150–2156 (2007).

79. Falótico, T., Valença, T., Verderane, M. P. & Fogaça, M. D. Stone tools differences across three capuchin monkey populations: food’s physical properties, ecology, and culture. Sci. Rep. 12, 14365 (2022).

80. Proffitt, T., Reeves, Jonathan. S., Pacome, S. S. & Luncz, Lydia. V. Identifying functional and regional differences in chimpanzee stone tool technology. R. Soc. Open Sci. 9, 220826 (2022).

81. Sakura, O. & Matsuzawa, T. Flexibility of Wild Chimpanzee Nut-cracking Behavior Using Stone Hammers and Anvils: an Experimental Analysis. Ethology 87, 237–248 (1991).

82. Neufuss, J., Humle, T., Cremaschi, A. & Kivell, T. L. Nut-cracking behaviour in wild-born, rehabilitated bonobos (*Pan paniscus*): a comprehensive study of hand-preference, hand grips and efficiency. Am. J. Primatol. 79, e22589 (2016).

83. Malherbe, M. et al. Protracted development of stick tool use skills extends into adulthood in wild western chimpanzees. PLoS Biol. 22, e3002609 (2024).

84. Schuppli, C. et al. Observational social learning and socially induced practice of routine skills in immature wild orang-utans. Anim. Behav. 119, 87–98 (2016).

85. Berdugo, S., Cohen, E., Davis, A. J., Matsuzawa, T. & Carvalho, S. The ontogeny of chimpanzee technological efficiency. Preprint at 10.1101/2024.07.31.605100 (2024).

86. Lindenberger, U. Human cognitive aging: Corriger la fortune? Science 346, 572–578 (2014).

87. Paccagnella, M. Age, Ageing and Skills: Results from the Survey of Adult Skills. vol. 132 https://www.oecd-ilibrary.org/education/age-ageing-and-skills_5jm0q1n38lvc-en (2016).

88. Blesa, J., Trigo-Damas, I., del Rey, N. L.-G. & Obeso, J. A. The use of nonhuman primate models to understand processes in Parkinson’s disease. J. Neural. Transm. 125, 325–335 (2018).

89. Emborg, M. E. Nonhuman Primate Models of Neurodegenerative Disorders. ILAR J. 58, 190–201 (2017).

90. Arnsten, A. F. T. et al. Alzheimer’s-like pathology in aging rhesus macaques: Unique opportunity to study the etiology and treatment of Alzheimer’s disease. Proc. Natl. Acad. Sci. U.S.A. 116, 26230–26238 (2019).

91. Edler, M. K. et al. Aged chimpanzees exhibit pathologic hallmarks of Alzheimer’s disease. Neurobiol. Aging 59, 107–120 (2017).

92. Lemere, C. A. et al. Cerebral Amyloid-Beta Protein Accumulation with Aging in Cotton-Top Tamarins: A Model of Early Alzheimer’s Disease? Rejuv. Res. 11, 321–332 (2008).

93. King, B. J. Extractive foraging and the evolution of primate intelligence. Hum. Evol. 1, 361–372 (1986).

94. Estrada, A. et al. Impending extinction crisis of the world’s primates: Why primates matter. Sci. Adv. 3, e1600946 (2017).

95. Humle, T. The Tool Repertoire of Bossou Chimpanzees. in The Chimpanzees of Bossou and Nimba (eds. Matsuzawa, T., Humle, T. & Sugiyama, Y.) 61–72 (Springer Japan, Tokyo, Japan, 2011).

96. Bain, M. et al. Automated audiovisual behavior recognition in wild primates. Sci. Adv. 7, eabi4883 (2021).

97. Friard, O. & Gamba, M. BORIS: a free, versatile open-source event-logging software for video/audio coding and live observations. Methods Ecol. Evol. 7, 1325–1330 (2016).

98. Boesch, C. et al. Chimpanzee ethnography reveals unexpected cultural diversity. *Nat*. Hum. Behav. 4, 910–916 (2020).

99. Boesch, C., Bombjaková, D., Meier, A. & Mundry, R. Learning curves and teaching when acquiring nut-cracking in humans and chimpanzees. Sci. Rep. 9, 1515 (2019).

100. Bates, D., Mächler, M., Bolker, B. & Walker, S. Fitting Linear Mixed-Effects Models Using **lme4**. J. Stat. Soft. 67, (2015).

