## Supplementary_Materials for "Old age variably impacts chimpanzee engagement and efficiency in stone tool use"

**Table S1. Ethogram of codable manipulations for observations of nut-cracking behaviors*.** Manipulations in bold are coded alongside a corresponding object. There are 6 possible corresponding objects: 1. Nut, 2. Hammer, 3. Anvil, 4. Kernel, 5. Shell, 6. Bare Hand. Codes in italics are used to denote the start and end of observable sequences. Coding commenced when individuals began interacting with stones, nuts, or nut-fragments. Coding ceased when individuals began engaging in a new behavior e.g. play, grooming, eating oil-palm fruits. On the occasions where an individual moved out of clear sight of the video recording, or behavior became obscured by an individual's body position, sequences were terminated with a 'Not Visible' codon, and marked as incomplete. This ethogram has been used in previous studies to collect data from the Bossou archive, see manuscript reference Howard-Spink et al. 2024.

| Action | Description |
|---|---|
| **Bite** | Place object in mouth and bite with teeth. Differs from 'Eat' as there is no consumption. Differs from 'Store' as object moves in and out of mouth whilst chimp is stationary. Differs from 'Peel Teeth' as bite applies general force, whereas peel with mouth is to remove shell fragments from kernel dexterously. Differs from 'Kiss' as object enters inside the mouth. |
| **Brush** | Brush objects off the anvil. |
| **Drop** | Place object(s) on ground. |
| **Eat** | Consume object. Differs from bite and store as it requires successful ingestion. |
| **Flip** | Flip object over. |
| **Grasp** | Grasp an object and move off the ground. |
| **Kick** | Apply rapid, hard force on object with the foot, so that the object is displaced. |
| **Kiss** | Place object to lips or nose, but not inside of the mouth. |
| **Pass** | Pass object between hands. |
| **Peel Hand** | Peel shell off of kernel with hands. |
| **Peel Teeth** | Peel shell off of kernel with teeth. |
| **Place** | Place an object on an anvil. |
| **Provide** | Directly hand an object to another individual. |
| **Pull Foot** | Pull an object across the ground with foot. |
| **Pull Hand** | Pull an object across the ground with hand. |
| **Push Foot** | Push an object across the ground with foot. |
| **Push Hand** | Push an object across the ground with hand. |
| **Rake Foot** | Pull many objects towards oneself using foot/leg. |
| **Rake Hand** | Pull many objects towards oneself using hand/arm. |
| **Reorient** | Rotate object horizontally. |
| **Roll Foot** | Roll object along the floor with foot. |
| **Roll Hand** | Roll object along the floor with hand. |
| **Spit** | Let object fall from mouth or lips. |
| **Stomp** | Whilst standing, apply strong force to object with foot. |
| **Store** | Place object(s) in mouth for transportation. Separated from bite, eat, peel teeth as it is followed by transportation across the outdoor laboratory, before then being removed from the mouth intact. |
| **Strike One Hand** | Strike with one hand, (and associated object). |
| **Strike Two Hand** | Strike with two hands, (and associated object). |
| **Support Foot** | Support object with foot. |
| **Support Hand** | Support object with hand. |
| **Take** | Receive an object directly from another individual. |
| **Throw** | Throw object away from self, horizontally or vertically. |
| **Touch Foot** | Touch or grasp object without moving it using foot. |
| **Touch Hand** | Touch or grasp object without moving it using hand. |
| Relocate | Stand up and move to a new area; is immediately followed by another coding action. |
| *Start* | Start of a sequence. |
| *End Bout* | End of a sequence. |
| *Not Visible* | Individual moves out of view, and coding cannot continue. Differs from 'End Bout', as there is no evidence the individual has stopped engaging with stones, nuts or nut-fragments. |

**Table S2. Model outputs for attendance rates over each year.**

glmer(Encounters ~ Year-Scaled*Age_Cohort + (1| ID) + offset(log(Field_Season_Duration)), family = Poisson)

| Random Effect | Variance | SD | | |
|---|---|---|---|---|
| ID (Intercept) | 0.055 | 0.235 | | |
| Fixed Effect | Estimate | SE | z value | p |
| (Intercept) | -0.570 | 0.107 | -5.35 | <0.001 |
| Year-Scaled | -0.123 | 0.078 | -1.57 | 0.116 |
| Old_Cohort | -0.198 | 0.141 | -1.40 | 0.160 |
| Year-Scaled:Old_Cohort | -0.220 | 0.096 | -2.29 | 0.022 |

**Table S3. The number of stone tool selection events sampled for each individual in each field season.**

| Individual | Year | | | | | Total |
|---|---|---|---|---|---|---|
| | 1999 | 2004 | 2008 | 2011 | 2016 | |
| Fana | 2 | 2 | 8 | 0 | 4 | 16 |
| Jire | 4 | 4 | 5 | 3 | 3 | 19 |
| TUA | 6 | 9 | 7 | 3 | 0 | 25 |
| Velu | 3 | 5 | 10 | 0 | 3 | 21 |
| Yo | 4 | 5 | 8 | 9 | 1 | 27 |

**Table S4. Models of stone-tool selection duration across years, with AIC**. AIC values reported for models fitted by Maximum Likelihood. PAB is the encounter in which stone tool selection events occurred.

| Model | Formula | df | AIC |
|---|---|---|---|
| REML = F | | | |
| Individual Random Slope and Intercept for Year-Scaled. | Log(Time) ~ Year-Scaled + Stones Selected + Stones Previously Taken + (1 + Year-Scaled\|\|ID) + (1\|PAB) | 8 | 231 |
| Dropped Year-Scaled | Log(Time) ~ Stones Selected + Stones Previously Taken + (1\|ID) + (1\|PAB) | 6 | 234 |
| Null for All Fixed Effects | Log(Time) ~ 1 + (1\|ID)+ (1\|PAB) | 4 | 234 |
| REML = T | | | |
| Random Slope & Intercept | Log(Time) ~ Year-Scaled + Stones Selected + Stones Previously Taken + (1 + Year-Scaled\|\|ID) + (1\|PAB) | 8 | 244 |
| Fixed Slope & Random Slope | Log(Time) ~ Year-Scaled + Stones Selected + Stones Previously Taken + (1\|ID) + (1\|PAB) | 7 | 246 |

**Table S5. Summary output for mixed-effect model with individual as a random slope, and encounter and individual as random intercepts.** 108 observations, 49 encounters, 5 individuals.

Log(Time) ~ Year-Scaled + Stones Selected + Stones Previously Taken + (1 + Year-Scaled||ID) + (1|PAB), REML = F

| Random Effect | Variance | SD | |
|---|---|---|---|
| Encounter | 0.02 | 0.15 | |
| ID (Intercept) | 0.13 | 0.35 | |
| ID (Slope – Year-Scaled) | 0.05 | 0.22 | |
| (Residual) | 0.35 | 0.59 | |
| **Fixed Effect** | **Estimate** | **SE** | **t value** |
| (Intercept) | 1.86 | 0.22 | 8.43 |
| Year-Scaled | 0.14 | 0.12 | 1.18 |
| Two Stones Selected | 0.31 | 0.15 | 2.14 |
| Stones Previously Taken | -0.01 | 0.02 | -0.65 |

**Table S6. Individual coefficients for the mixed-effect model for tool selection duration, with individual as a random slope, and encounter and individual as random intercepts.**

| Individual | Intercept | Scaled Year (Slope) |
|---|---|---|
| Fana | 1.93 | 0.27 |
| Jire | 1.80 | 0.17 |
| TUA | 1.39 | -0.17 |
| Velu | 1.76 | 0.07 |
| Yo | 2.41 | 0.35 |

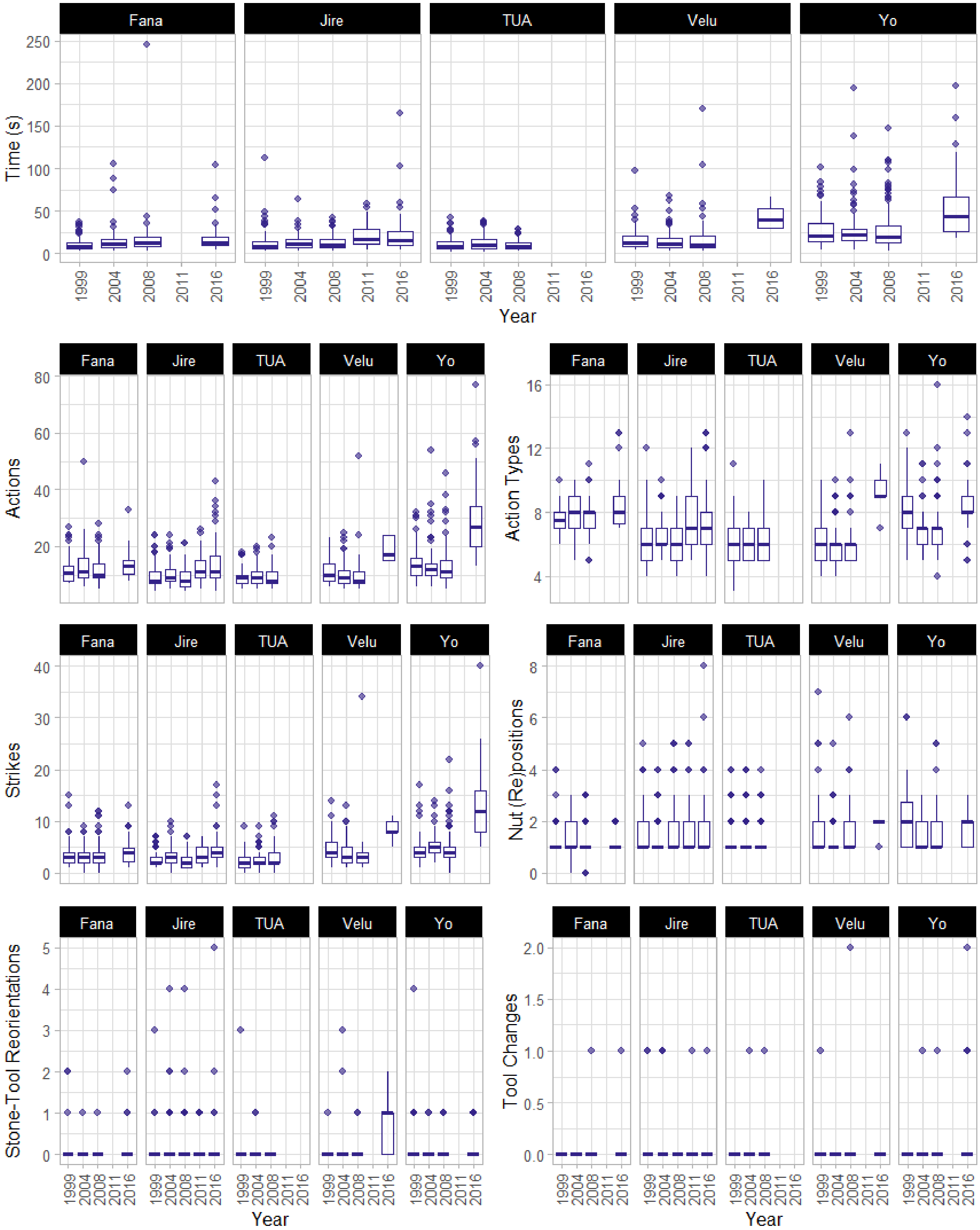


**Figure S1. Boxplots describing metrics of efficiency for oil-palm nut cracking and processing, for each individual in each field season.**

**Table S7. Summary output for the random slope mixed-effect model for the total time taken to crack and process oil-palm nuts over sampled field seasons.** 1538 observations, 20 encounters, 5 individuals. Values rounded to 3.d.p.

lmer(Log_Time ~ Year-Scaled + (Year-Scaled | ID) + (1| Encounter) , REML = F)

| Random Effect | Variance | SD | |
|---|---|---|---|
| Encounter | 0.029 | 0.169 | |
| ID (Intercept) | 0.089 | 0.298 | |
| ID (Slope – Year-Scaled) | 0.002 | 0.039 | |
| (Residual) | 0.418 | 0.647 | |
| **Fixed Effect** | **Estimate** | **SE** | **t value** |
| (Intercept) | 2.657 | 0.142 | 18.767 |
| Year-Scaled | 0.077 | 0.030 | 2.605 |

**Table S8. Summary output for the random slope mixed-effect model for the number of discrete actions used to crack and process oil-palm nuts over sampled field seasons.** 1538 observations, 20 encounters, 5 individuals. Values rounded to 3.d.p.

glmer(Action Events ~ Scaled Year + (Year-Scaled |ID) + (1|Encounter), family = poisson)

| Random Effect | Variance | SD | |
|---|---|---|---|
| Encounter | 0.084 | 0.289 | |
| ID (Intercept) | 0.029 | 0.170 | |
| ID (Slope – Year-Scaled) | 0.003 | 0.059 | |
| Fixed Effect | Estimate | SE | z value |
| (Intercept) | 2.563 | 0.102 | 25.241 |
| Year-Scaled | 0.022 | 0.029 | 0.769 |

**Table S9. Summary output for the random slope mixed-effect model for the number strikes of the hammer stone used to crack and process oil-palm nuts over sampled field seasons.** 1538 observations, 20 encounters, 5 individuals. Values rounded to 3.d.p.

glmer(Strikes ~  Year-Scaled + ( Year-Scaled |ID) + (1|Encounter), family = poisson)

| Random Effect | Variance | SD | |
|---|---|---|---|
| Encounter | 0.157 | 0.397 | |
| ID (Intercept) | 0.042 | 0.204 | |
| ID (Slope – Year-Scaled) | 0.011 | 0.104 |  |
| Fixed Effect | Estimate | SE | z value |
| (Intercept) | 1.462 | 0.131 | 11.149 |
| Scaled Year | 0.079 | 0.051 | 1.545 |

**Table S10. Intercept and slope for individual random effects for nut metrics of total time, total number of actions, and total number of strikes.** Values rounded to 3.d.p.

| Metric | Individual | Intercept | Year-Scaled (Slope) |
|---|---|---|---|
| **Total Time** | **Fana** | 2.614 | 0.064 |
| | **Jire** | 2.508 | 0.089 |
| | **TUA** | 2.328 | 0.036 |
| | **Velu** | 2.635 | 0.068 |
| | **Yo** | 3.200 | 0.128 |
| **Actions** | **Fana** | 2.685 | -0.028 |
| | **Jire** | 2.507 | 0.049 |
| | **TUA** | 2.416 | 0.043 |
| | **Velu** | 2.399 | -0.050 |
| | **Yo** | 2.814 | 0.098 |
| **Strikes** | **Fana** | 1.415 | 0.008 |
| | **Jire** | 1.321 | 0.051 |
| | **TUA** | 1.290 | 0.195 |
| | **Velu** | 1.476 | -0.043 |
| | **Yo** | 1.820 | 0.186 |

**Table S11. Summary of nut cracking and processing metrics for Tua, Jire and Yo between 2008 and 2016.** Total time duration is summarized by mean and standard deviation. All other metrics are summarized by medians and interquartile ranges. Horizontal lines indicate where no data is available for a given year. Information on how each metric is defined and estimated can be found in the Methods.

| Individual | Metric | Year and Nut Type | | | |
|---|---|---|---|---|---|
| | | 2008 | 2011 | | 2016 |
| **TUA** | | **oil-palm (n = 81)** | **coula (n = 26)** | | **-** |
| | *Means (sd)* | | | | |
| | Total Time (s) | 10.6 (6.34) | 54.7 (92.6) | | - |
| | *Medians (IQR)* | | | | |
| | Actions | 8 (4) | 25.5 (16) | | - |
| | Action Types | 6 (2) | 8 (3) | | - |
| | Strikes | 2 (2) | 7 (6.75) | | - |
| | Nut Positioning | 1 (0) | 2 (2.75) | | - |
| | Tool Reorientations | 0 (0) | 0 (0.75) | | - |
| | Tool Changes | 0 (0) | 0 (0) | | - |
| **Jire** | | **oil-palm (n = 104)** | **oil-palm (n = 25)** | **coula (n = 17)** | **oil-palm (n = 82)** |
| | *Means (sd)* | | | | |
| | Total Time (s) | 12.8 (8.37) | 21.8 (14.9) | 44.6 (21.6) | 21.9 (22.3) |
| | *Medians (IQR)* | | | | |
| | Actions | 8 (5) | 11 (6) | 26 (17) | 11 (7.75) |
| | Action Types | 6 (2) | 7 (3) | 11 (2) | 7 (2) |
| | Strikes | 2 (2) | 3 (3) | 7 (4) | 4 (2) |
| | Nut Positioning | 1 (1) | 1 (1) | 3 (1) | 1 (1) |
| | Tool Reorientations | 0 (0) | 0 (0) | 0 (1) | 0 (0) |
| | Tool Changes | 0 (0) | 0 (0) | 0 (0) | 0 (0) |
| **Yo** | | **oil-palm (n = 141)** | **coula (n = 20)** | | **oil-palm (n = 33)** |
| | *Means (sd)* | | | | |
| | Total Time (s) | 28.1 (24.6) | 186 (183.4) | | 55.9 (42.3) |
| | *Medians (IQR)* | | | | |
| | Actions | 11 (6) | 88.5 (60.5) | | 27 (14) |
| | Action Types | 7 (1) | 11.5 (8.5) | | 8 (1) |
| | Strikes | 4 (2) | 27 (23) | | 12 (8) |
| | Nut Positioning | 1 (1) | 11 (22.25) | | 2 (1) |
| | Tool Reorientations | 0 (0) | 0.5 (4.25) | | 0 (0) |
| | Tool Changes | 0 (0) | 0 (7.25) | | 0 (0) |

### Coula Nut Cracking

#### Results

Three of our focal individuals were observed cracking coula nuts in 2011 (Jire, Tua & Yo). Coula nuts do not occur naturally at Bossou, but are cracked by chimpanzees at other sites across Africa[1,2] and have historically been experimentally provided to individuals at Bossou during selected field seasons[2,3]. Of the three individuals who were observed cracking coula nuts in 2011, only Jire was observed also cracking oil-palm nuts in the same year. Coula nuts require somewhat more effort to crack and process than smaller oil-palm nuts[1]; however, for two individuals (Jire and Tua), coula nuts were cracked and processed with comparable efficiency to smaller oil-palm nuts (with a slight increase in total time taken, the number of actions and unique action types used, and the total number of strikes of the hammer stone; see Fig. S2 and Table S11 for corresponding data). On average, Jire took an extra 22.8 s to crack coula nuts compared with oil-palm nuts in 2011. Similarly, Tua took an additional 44.1s to crack coula nuts in 2011, compared with oil-palm nuts in 2008 (the closest comparison point for Tua, who was not observed cracking oil-palm nuts in 2011).

Conversely, Yo showed a considerably lower efficiency cracking coula nuts in 2011 compared with her efficiency cracking oil-palm nuts in the previously sampled field season (2008; see Fig. S2). On average, Yo took an additional 2.63 minutes (+157.9 s; mean duration across oil-palm nuts in 2008 and coula nuts in 2011) to crack open coula nuts, during which Yo performed an average of 78 additional actions to crack the nut and consume all associated kernel (including an additional 23 hammer strikes; measured as using median values for each nut species). For Yo, coula nuts frequently rolled off of the anvil stone, resulting in Yo placing each coula nut on the anvil 11 times prior to successful cracking (median value; in 2008, Yo would place oil-palm nuts on the anvil once per nut). Additionally, Yo exhibited an exceptional variability between the number of times she switched out tools during coula-nut cracking (interquartile range = 7.25, for oil-palm nuts in 2008 this was 0), as well as for the number of tool adjustments per coula nut (coula = 4.25, oil-palm = 0). Overall, these metrics suggest that Yo found coula-nut cracking in 2011 disproportionately difficult compared with oil-palm nut cracking. This was to an extent which was not mirrored by other individuals of similar ages, as well as compared to her own performance when cracking oil-palm nuts in previous and subsequent years (see Table S11).

#### Discussion

The disproportionate difficulty that Yo faced cracking Coula nuts was surprising given the history of this behavior at Bossou[2–4]. Unlike oil-palm nuts, whose trees occur within the home range of the Bossou community, coula nuts are not found at Bossou[2,3]. Coula nuts were first introduced to chimpanzees at the outdoor laboratory in 1993, when Yo was the only adult individual to begin cracking coula nuts without engaging in a phase of exploratory behaviour[2]. Other individuals began cracking coula nuts through successive, intermittent presentations of coula nuts across further field seasons (1996, 2000, 2002, 2005). Given that Yo readily engaged in coula nut cracking, it was deduced that Yo was likely an immigrant from the neighboring Yealé population, where both oil-palm and coula nuts are readily available[2,3]. Therefore, Yo likely has the longest history, and greatest experience cracking coula nuts, despite exhibiting such low efficiency in 2011.

There are numerous hypotheses for why Yo showed a dramatic reduction in efficiency when cracking coula nuts in 2011. It is important to note that, for Yo, data for the cracking of coula nuts in this year came from a single, three-hour long encounter, where Yo spent one hour and 17 minutes of continuous effort cracking the first 20 coula nuts. It is possible that a short-term effect (such as dehydration, hunger, physical injury) may have inhibited the cognitive processes or physical movements required for streamlined tool action. However, we noticed no obvious signs of physical injury, and Yo spent the majority of the encounter prioritizing nut cracking over both the water point and the more readily available oil-palm fruits. This reduces the likelihood of these hypotheses explaining the observed changes in Yo's behavior. Alternatively, it may be that the reduced ability to crack coula nuts in 2011 was a product of senescence, a hypothesis which is supported by the fact that Yo also exhibited the largest reduction in efficiency when cracking oil-palm nuts at older ages (which continued to intensify further in later life, past the 2016 field season; see below). Given that coula nuts are larger and rounder than oil-palm nuts (and thus can be more liable to roll off of anvil stones) and have more fibrous outer shells[1], they therefore may require heavier hammer stones, and/or more forceful or elaborate sequences of striking and peeling actions to separate shell from kernel. Physiological senescence – such as reduced strength of dexterity - may have rendered coula nut cracking even more challenging than the cracking of oil-palm nuts, for which Yo was also exhibiting increasing difficulty. Alternatively, cognitive senescence may have also contributed for the additional difficulty Yo faced when cracking coula nuts, where she may have struggled to generate suitable action patterns to crack open a less frequently encountered nut species.

Given the absence of data for coula nut cracking in earlier years, we emphasize that these conclusions are tentative, and further data collection is required for coula-nut cracking over different field seasons.

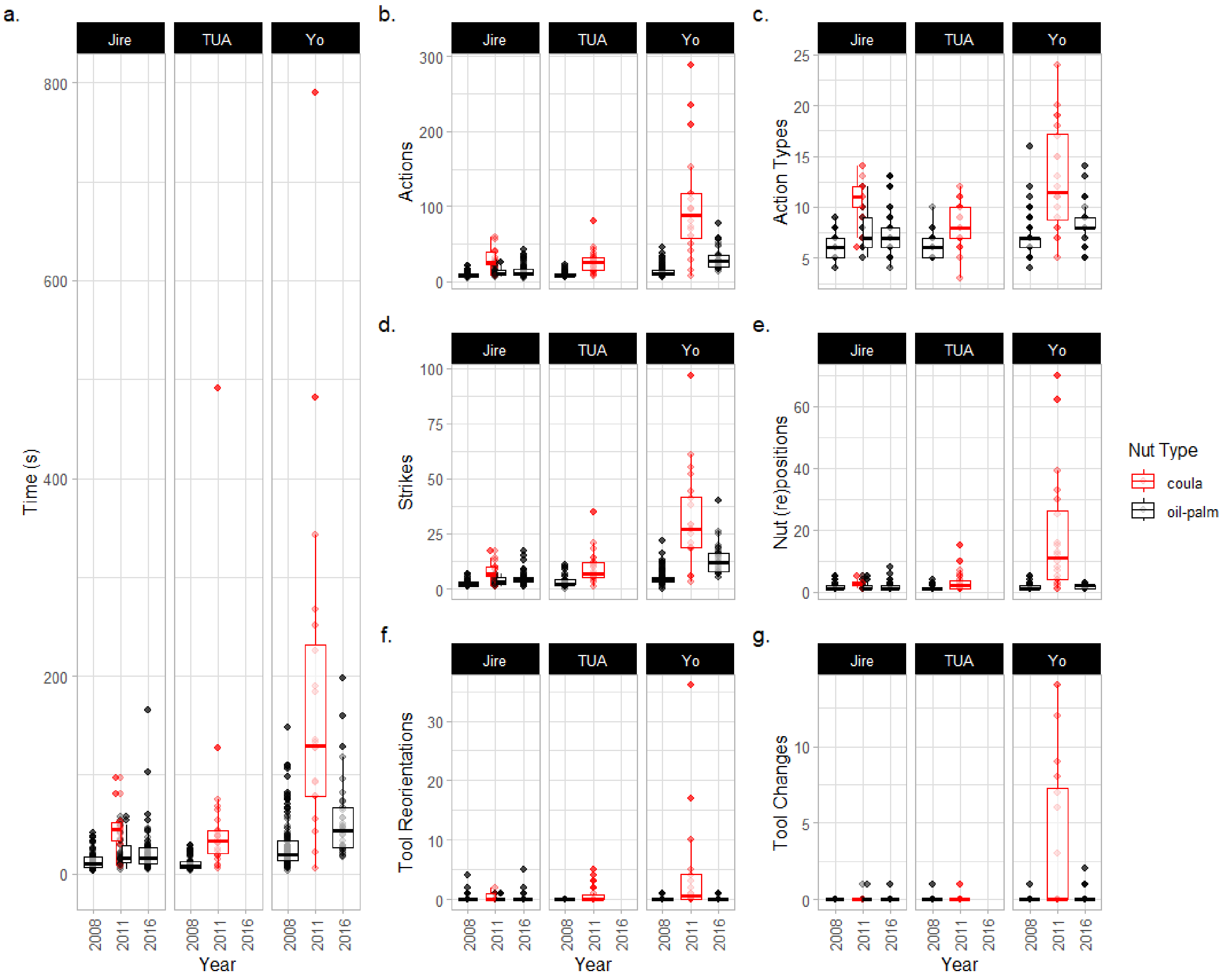


**Figure S2. Metrics for the cracking and processing of both oil-palm and coula nuts.** Data is confined to individuals who cracked nuts from both species. Data for coula nuts is in red, and data for oil-palm nuts is in black. Data describes the cracking and processing of individual nuts, including (a) the total time taken; (b) the total number of actions used; (c) the number of unique types of actions used; (d) the number of hammer strikes; (e) the number of times the nut had to be placed and replaced on the anvil; (f) the number of reorientations of stone tools, and (g) the number of times stone tools were switched over, or switched out for new tools.

**Table S12.** Each unique action type employed by Yo in 1999 and 2016, described as a proportion of the first 1000 actions observed for Yo in each year. The difference between years is found, and whether this difference is greater than 0.05 (5%) is determined. A direction of change between years is also indicated for each action.

| Action | 1999 | 2016 | 2016 - 1999 | Difference > 0.05? | direction |
|---|---|---|---|---|---|
| brush SHELL | 0.01632302 | 0.00981354 | 0.00650948 | FALSE | decrease |
| drop ANVIL | 0.00429553 | 0.00392542 | 0.00037012 | FALSE | decrease |
| drop HAMMER | 0.03006873 | 0.00785083 | 0.02221789 | FALSE | decrease |
| drop KERNEL | 0.00085911 | 0.00294406 | 0.00208496 | FALSE | increase |
| drop NUT | 0.00429553 | 0.00392542 | 0.00037012 | FALSE | decrease |
| drop SHELL | 0.01546392 | 0.02551521 | 0.01005129 | FALSE | increase |
| eat KERNEL | 0.08676976 | 0.13837095 | 0.05160119 | TRUE | increase |
| flip ANVIL | 0.00257732 | 0 | 0.00257732 | FALSE | decrease |
| flip HAMMER | 0.00085911 | 0 | 0.00085911 | FALSE | decrease |
| flip KERNEL | 0.00085911 | 0 | 0.00085911 | FALSE | decrease |
| grasp ANVIL | 0.00429553 | 0.00392542 | 0.00037012 | FALSE | decrease |
| grasp HAMMER | 0.04553265 | 0.00588813 | 0.03964452 | FALSE | decrease |
| grasp KERNEL | 0.07044674 | 0.05888126 | 0.01156548 | FALSE | decrease |
| grasp NUT | 0.11340206 | 0.05691855 | 0.05648351 | TRUE | decrease |
| kiss KERNEL | 0.00257732 | 0 | 0.00257732 | FALSE | decrease |
| kiss NUT | 0.00085911 | 0 | 0.00085911 | FALSE | decrease |
| pass HAMMER | 0.00085911 | 0 | 0.00085911 | FALSE | decrease |
| pass KERNEL | 0.00171821 | 0 | 0.00171821 | FALSE | decrease |
| pass NUT | 0.01718213 | 0.00196271 | 0.01521942 | FALSE | decrease |
| peelhand SHELL | 0.00257732 | 0.00392542 | 0.0013481 | FALSE | increase |
| peelteeth SHELL | 0.05584192 | 0.12365064 | 0.06780871 | TRUE | increase |
| place HAMMER | 0.01202749 | 0.00196271 | 0.01006478 | FALSE | decrease |
| place KERNEL | 0.00601375 | 0.00098135 | 0.00503239 | FALSE | decrease |
| place NUT | 0.09621993 | 0.0451423 | 0.05107763 | TRUE | decrease |
| relocate | 0.00687285 | 0.00294406 | 0.00392879 | FALSE | decrease |
| reorient ANVIL | 0.00945017 | 0.00392542 | 0.00552475 | FALSE | decrease |
| rollhand HAMMER | 0.00085911 | 0 | 0.00085911 | FALSE | decrease |
| spit SHELL | 0.00085911 | 0.00098135 | 0.00012225 | FALSE | increase |
| strikeonehand HAMMER | 0.35395189 | 0.44455348 | 0.09060159 | TRUE | increase |
| strikeonehand NUT | 0.00085911 | 0 | 0.00085911 | FALSE | decrease |
| supportfoot ANVIL | 0.00343643 | 0.00294406 | 0.00049236 | FALSE | decrease |
| touchfoot ANVIL | 0.00085911 | 0 | 0.00085911 | FALSE | decrease |
| touchhand ANVIL | 0.00429553 | 0.00392542 | 0.00037012 | FALSE | decrease |
| touchhand HAMMER | 0.00515464 | 0 | 0.00515464 | FALSE | decrease |
| touchhand KERNEL | 0.00601375 | 0.00294406 | 0.00306968 | FALSE | decrease |
| touchhand NUT | 0.01460481 | 0.03140334 | 0.01679853 | FALSE | increase |
| touchhand SHELL | 0.00085911 | 0.00392542 | 0.00306631 | FALSE | increase |
| bite SHELL | 0 | 0.00098135 | 0.00098135 | FALSE | increase |
| brush HAMMER | 0 | 0.00098135 | 0.00098135 | FALSE | increase |
| grasp SHELL | 0 | 0.00294406 | 0.00294406 | FALSE | increase |
| reorient HAMMER | 0 | 0.00098135 | 0.00098135 | FALSE | increase |
| supporthand ANVIL | 0 | 0.00098135 | 0.00098135 | FALSE | increase |

#### 2018 Data: Yo Cracking Oil-Palm Nuts

Data collection at the outdoor laboratory (second location) was conducted in 2018, offering an additional year for our analysis. We did not, however, include this data in our main manuscript for several reasons.

Data was collected using camera traps in 2018, which were motion triggered. These camera traps were only able to capture a limited viewpoint of the outdoor laboratory, and did not record encounters continuously. Therefore, estimating the entire party composition during encounters; the behaviors of individuals as they moved around the outdoor laboratory, and the duration of stone-tool selection events were not possible, as individuals were often out of sight for some or all of the behavior. We also could not collect fine-grained action sequence data for the entire behavioral sequence used to crack individual nuts, as there were gaps in the video footage lasting several seconds. Additionally, in 2018, only three elderly individuals remained at Bossou (Yo, Jire & Fana), and the majority of clear video footage concerning nut cracking was of Yo. All nuts cracked by Yo in this year were oil-palm nuts.

Whilst the collection of fine-grained action sequence data was not possible, it was possible to record the start and end times of nut cracking sequences aimed at individual nuts using timestamps on video footage. We therefore collected the start and end times of nut cracking sequences performed by Yo. We filtered our data to only include nuts where we could clearly identify the start and end of Yo's nut cracking behaviors across videos, using the same coding scheme reported in the methods of our manuscript (see Table S13).

To calculate a mean duration of oil-palm nut cracking for Yo in 2018, we first found the mean within each encounter, and then calculated a mean across encounters. This blocking was used to compensate for nonindependence of nuts cracked as part of the same encounter. Overall, it took Yo on average 74.2 s to crack each oil palm nut in 2018 (SD = 9.1 s; estimated using the means of each encounter). This was 18.3 s longer than the mean time taken in 2016, meaning that Yo became even less efficient at oil-palm nut cracking in later years. Overall, when comparing the mean duration of oil-palm nut cracking between 1999 and 2018, Yo experienced an +171% change in total time taken (1999 = 27.4 s; difference between years = +46.7 s).

**Table S13. Total time taken for Yo to crack oil-palm nuts in 2018**. Encounter numbers indicate where data was collected for the same encounter. Nut number indicates which nut (in chronological order) Yo was cracking for a specific encounter. Numbers missing from this chronology were removed as we could not reliably estimate start and end times.

| Encounter | Nut Number | Duration (s) |
|---|---|---|
| 9 | 6 | 145 |
| 9 | 7 | 73 |
| 9 | 8 | 24 |
| 9 | 9 | 29 |
| 11 | 1 | 220 |
| 11 | 3 | 173 |
| 11 | 4 | 165 |
| 11 | 5 | 54 |
| 11 | 6 | 42 |
| 11 | 7 | 17 |
| 11 | 8 | 36 |
| 11 | 9 | 75 |
| 11 | 10 | 37 |
| 11 | 17 | 35 |
| 11 | 18 | 59 |
| 11 | 19 | 54 |

#### Supplementary References

1. Boesch, C. & Boesch, H. Optimisation of Nut-Cracking With Natural Hammers By Wild Chimpanzees. *Behav.* **83**, 265–286 (1983).
2. Biro, D., Sousa, C. & Matsuzawa, T. Ontogeny and cultural propagation of tool use by wild chimpanzees at Bossou, Guinea: Case studies in nut cracking and leaf folding. in *Cognitive Development in Chimpanzees* (eds. Matsuzawa, T., Tomonaga, M. & Masayuki, T.) 476–508 (Springer Tokyo, Tokyo, Japan, 2006).
3. Biro, D. *et al.* Cultural innovation and transmission of tool use in wild chimpanzees: evidence from field experiments. *Anim. Cogn.* **6**, 213–223 (2003).
4. Matsuzawa, T. Field Experiments on use of Stone Tools in the wild. in *Chimpanzee Cultures* (eds. Wrangham, R. W., McGrew, W. C., De Waal, F. B. M. & Heltne, P. G.) 351–370 (Harvard University Press, 1994).
